# RESP2: An uncertainty aware multi-target multi-property optimization AI pipeline for antibody discovery

**DOI:** 10.1101/2024.07.30.605700

**Authors:** Jonathan Parkinson, Ryan Hard, Young Su Ko, Wei Wang

**Affiliations:** Department of Chemistry and Biochemistry, University of California, San Diego, La Jolla, CA 92093-0359; Department of Cellular and Molecular Medicine, University of California, San Diego, La Jolla, CA 92093-0359; MAP Bioscience, La Jolla, CA 92093

## Abstract

Discovery of therapeutic antibodies against infectious disease pathogens presents distinct challenges. Ideal candidates must possess not only the properties required for any therapeutic antibody (e.g. specificity, low immunogenicity) but also high affinity to many mutants of the target antigen. Here we present RESP2, an enhanced version of our RESP pipeline, designed for the discovery of antibodies against one or multiple antigens with simultaneously optimized developability properties. We first evaluate this pipeline *in silico* using the Absolut! database of scores for antibodies docked to target antigens. We show that RESP2 consistently identifies sequences that bind more tightly to a group of target antigens than any sequence present in the training set with success rates >= 85%. Popular generative AI techniques evaluated on the same datasets achieve success rates of 1.5% or less by comparison. Next we use the receptor binding domain (RBD) of the COVID-19 spike protein as a case study, and discover a highly human antibody with broad (mid to high-affinity) binding to at least 8 different variants of the RBD. These results illustrate the advantages of this pipeline for antibody discovery against a challenging target. A Python package that enables users to utilize the RESP pipeline on their own targets is available at https://github.com/Wang-lab-UCSD/RESP2, together with code needed to reproduce the experiments in this paper.

## INTRODUCTION

Discovery of therapeutic antibodies against any fast mutating antigens such as those for infectious diseases and some cancers poses a unique array of challenges. For an antibody to be successful, it must not only possess good “developability” (e.g. minimal immunogenicity risk, robust solubility and stability)^1,2^, but also exhibit tight binding to multiple antigen variants to achieve broad neutralization against potential mutants.

Traditionally this multi-parameter optimization problem has been tackled through time-consuming experimental approaches^3–5^. Recently, a variety of machine learning (ML)-assisted approaches to expediting this process via predicting affinity or other key properties have emerged^6–13^. While ML models can learn complex relationships between structure and properties or between sequence and properties, their accuracy diminishes when applied to new data that deviate significantly from the training set^14–17^.

In our previous work, we introduced RESP, an ML-assisted pipeline for identifying antibodies with tight binding to a specific target^12^. This pipeline employs uncertainty-aware ML models that effectively manage the limits of their knowledge to avoid inappropriate extrapolation. Our algorithm also conducts *in silico* directed evolution to find tight binders to a target not necessarily present in the screening library.

In our current study, we enhance the RESP pipeline to enable *in silico* optimization against multiple antigens or antigen variants. We present two new uncertainty-aware ML models not previously used for protein engineering. We also integrate a generative model trained on 130 million human antibody sequences. This integration enables rapid assessment of humanness and developability for any antibody sequence suggested by the *in silico* evolution algorithm and humanization of promising candidates.

A variety of “zero-shot” generative AI approaches not requiring any experimental data have been proposed in recent work^18–22^. Despite this attractive advantage, currently available zero-shot methods suffer from highly variable success rates, in some cases identifying sequences with greatly improved binding and in other cases failing to improve binding significantly. The black-box nature of the models and the approach makes it difficult to determine in which cases it is most likely to succeed. Alternatively, inverse folding models predict sequence from structure in the hopes that modifying the sequence without changing the backbone structure may improve binding or some other property^21,23,24^. This approach too has achieved success rates which vary considerably by target and task, and a reliable starting complex structure may be needed to achieve good results. Finally, biophysics-based methods like MD simulations, FoldX scoring and free energy perturbation (FEP) are computationally expensive so that considerable computational resources are required, and the success rate achieved is still relatively modest.

Consequently, while zero-shot approaches have shown some promise, their drawbacks suggest that in scenarios where phage or yeast display capabilities are available, a machine learning-assisted approach such as the one we suggest here is highly competitive. As we will demonstrate, unlike either purely experimental or purely computational techniques, the RESP pipeline is able to achieve high success rates while simultaneously optimizing for developability.

We first assess the updated RESP pipeline’s efficacy on synthetic data using the Absolut! database and docking tool from Robert et al.^25^ The Absolut! software toolkit generates and scores lattice-based antibody-antigen binding structures. Since these structures are discretized, the software is able to dock all possible binding poses and score them using a physics-based scoring function, returning the best score obtained for each proposed CDRH3. The accompanying database contains precomputed scores for nearly 8 million CDRH3 sequence variants against 159 antigens. The authors demonstrate that rankings of ML model performance for different models on their synthetic dataset are well-correlated with rankings for the same model architectures on experimental antibody-antigen affinity data, suggesting that techniques which perform well here are likely to perform well on real-world data as well.

We extract five pairs and one triplet of antigens with > 50% pairwise percent identity from the Absolut! database and use either the RESP pipeline or two popular generative AI techniques to generate candidates against each target group. We demonstrate that RESP consistently identifies binders with higher affinity than any sequence present in the training set, achieving success rates >=85% in every case, while the generative AI techniques never achieve a success rate better than 1.5%.

We next demonstrate RESP2’s efficacy on real-world data using the receptor binding domain (RBD) of the SARS-CoV-2 spike protein as a target. The SARS-CoV-2 virus is the causative agent of the COVID-19 pandemic responsible for 7 million deaths worldwide^26^. Its spike protein is a challenging target for antibody drug discovery due to its mutability. Despite the approval of various antibody treatments for COVID-19, their effectiveness is often compromised by viral mutations and by the pathogen’s continuing evolution^27^. Beginning with an antibody exhibiting limited affinity across various RBDs, we employ RESP2 to identify, through a single additional round of screening and sequencing, highly human antibodies with an expanded range of affinities for multiple variants (e.g. BA.2) to which the original antibody had negligible affinity. These results are consistent with the high success rates observed for RESP in the *in silico* validation and suggest the RESP pipeline can be an efficient approach to multi-parameter optimization. Practitioners interested in utilizing the updated RESP pipeline for their own data and target can use the Python package available at https://github.com/Wang-lab-ucsd/RESP2.

## RESULTS

### Overview of the RESP2 pipeline

The RESP2 pipeline is enhanced from the previous version to facilitate discovery of antibodies that both bind to multiple target antigens and exhibit additional desired properties (Figure 1A).

**Figure 1.**
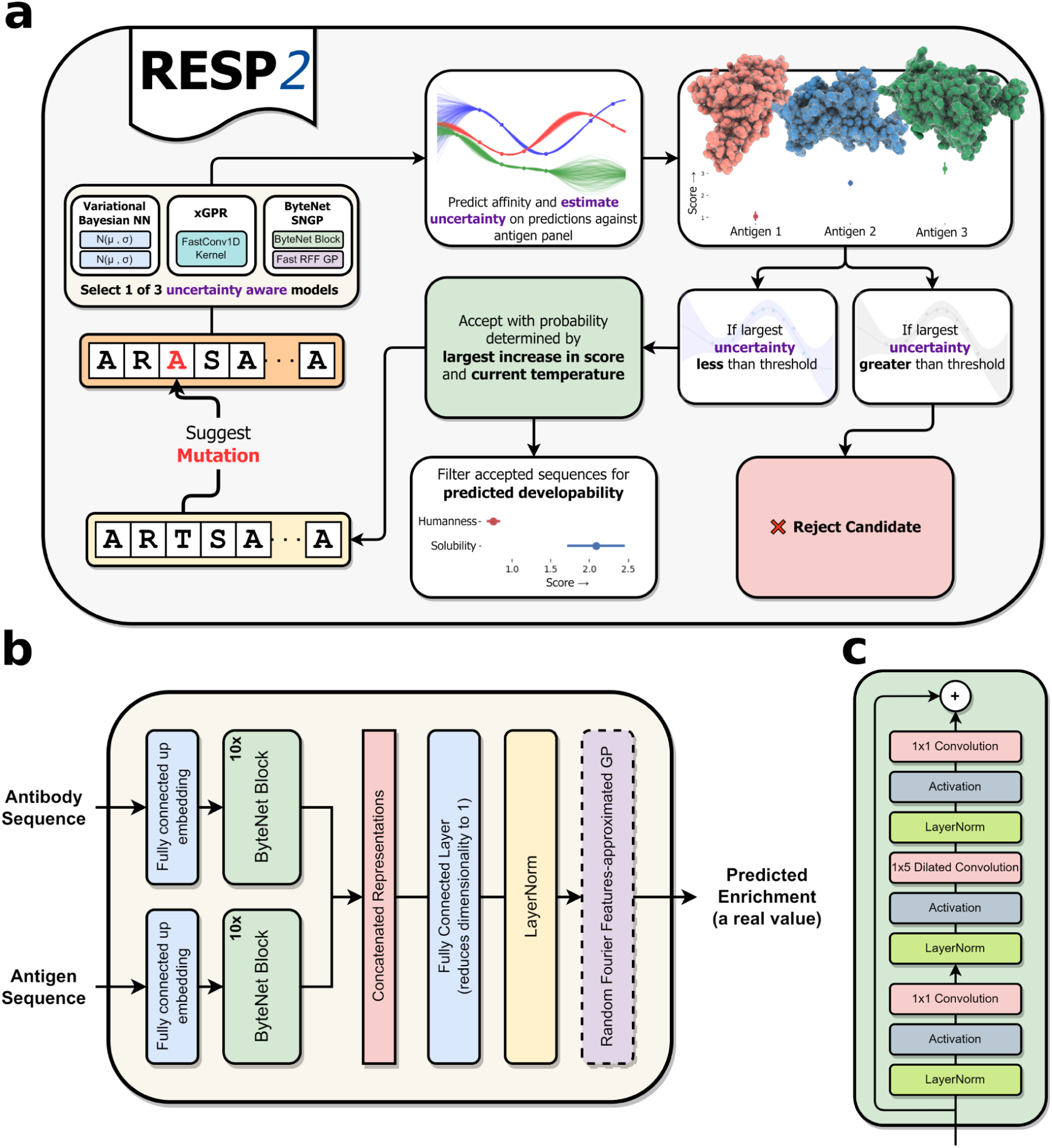
1A. An overview of the RESP2 pipeline. 1B. The overall ByteNet architecture. 1C. The ByteNet block used in the ByteNet architecture.

First, we employ random mutagenesis to generate a focused library of the scFv heavy chain. We screen this library against each antigen of interest and used fluorescence activated cell sorting (FACS) to select a subpopulation of mutants with improved binding to each antigen. Both the unscreened library and the tighter-binding subpopulations against each antigen are sequenced. Unlike RESP, RESP2 collects data for multiple antigens, facilitating concurrent optimization against multiple targets.

Second, we encode the raw data using a representation of the antibody and antigen sequences, then train an uncertainty-aware “affinity model” to predict the likelihood that a sequence is a strong binder. The affinity model takes a paired antibody-antigen sequence as input and predicts enrichment in a subpopulation as a proxy for binding affinity. RESP2 provides 3 methods to estimate uncertainty, including the variational Bayesian network used by RESP^12^, a fast and scalable approximate Gaussian process (GP) method implemented in xGPR^28^, and a new uncertainty-aware deep learning architecture (ByteNet-SNGP) developed in this work with the greatest flexibility for antibody engineering. We demonstrate that both xGPR and ByteNet-SNGP offer improved performance coupled with well-calibrated uncertainty quantitation and that uncertainty assigned by these models can identify out-of-distribution (OOD) sequences.

Third, we refine the *in silico* directed evolution (ISDE) algorithm we developed in RESP to concurrently optimize binding to multiple antigens. We begin with the wild-type as the initial candidate sequence. We then select positions for *in silico* mutagenesis using the affinity model from the previous step. During each iteration, the algorithm proposes a mutation to the current candidate. Mutations associated with high uncertainty in the model’s scoring are automatically discarded to avoid reliance on potentially inaccurate predictions. Next, the algorithm accepts or rejects the mutation in a simulated annealing process based on the smallest increase or largest decrease in score against any antigen rather than focusing on a single antigen as in RESP (see Materials and Methods for details). This procedure selects for mutations which enhance or at least preserve binding affinity against all the targets. The temperature in the simulated annealing is high at the start and gradually lowered to promote exploration in the early stages and refinement later on.

Finally, we collect the candidates generated by the ISDE and filter them to identify those with other desired properties in addition to high affinity for the antigens of interest. If this filtering process yields few or no candidate sequences, the ISDE algorithm is re-run to generate more candidates. We use SAM^29^, a fully interpretable generative model for distinguishing human from nonhuman antibodies, to score candidates to ensure low risk of immunogenicity.

In recent work, we^29^ performed an extensive validation of the SAM model on a series of benchmark datasets. Using a held-out test set of > 400 million human heavy and light chains, for example, we showed that SAM can distinguish human heavy and light chains from a diverse array of other species with accuracy and reliability superior to any other model in the literature (see Parkinson et al. for details). We also demonstrated that SAM achieves state-of-the art results for correlation between assigned scores and clinical outcomes (percent of patients forming anti-drug antibodies or ADA), so that sequences with improved SAM scores exhibit reduced risk of immunogenicity in the clinic. Unlike many existing alternatives, the SAM model is fully interpretable; it clusters its training data using a model which represents a distribution across sequence space. We can use this mixture model to score full sequences, specific framework or CDR regions or even specific positions in each sequence. In our previous work, we used the interpretable nature of the SAM tool to identify a previously unreported data quality issue with the Observed Antibody Space (OAS) database^30^. Given its demonstrated advantages, we use the SAM tool for scoring antibodies for humanness in this paper. We also score selected sequences for solubility. Additional filters can easily be added at this stage of the pipeline as desired.

### Validating the RESP pipeline on synthetic data

To further benchmark RESP2 against RESP1, to assess the usefulness of uncertainty and to compare with generative AI pipelines, we first conducted an *in silico* validation against thirteen antigens using the Absolut! software suite and database from Robert et al.^25^ The Absolut! software toolkit generates and scores lattice-based antibody-antigen binding structures. Since these structures are discretized, the software is able to dock all possible binding poses and score them using a physics-based scoring function, returning the best score obtained for each proposed CDRH3. The accompanying database contains precomputed scores for nearly 8 million CDRH3 sequence variants against 159 antigens. The authors demonstrate that rankings of ML model performance for different models on their synthetic dataset are well-correlated with rankings for the same model architectures on experimental antibody-antigen affinity data^25^. They suggest therefore that this toolkit can be useful for comparing model architectures and ML-assisted protein engineering strategies using relatively cheap *in silico* experiments that predict which strategy is likely to perform best on real data.

Since in this study we focus on design of broad-binding antibodies with high affinity to multiple mutated variants of an antigen, we first clustered the 159 antigens in the Absolut! Database using a 50% identity cutoff (see Materials and Methods, section “Experiments on synthetic data from the Absolut! Database” for details). We identified 5 pairs and 1 triplet where the sequences in the resulting cluster met our criteria. The percent identity for sequences in each target group ranges from 56% to 97.2% with an average of 78%. For each antigen, the Absolut! Database provides 500,000 CDRH3 sequences randomly sampled from the bottom 99% of the roughly 8 million CDRH3 evaluated against that antigen and the top 1% of CDRH3 sequences evaluated against that antigen, together with the docking score assigned by the Absolut! tool for each sequence.

We used only the bottom 500,000 sequence dataset (“weak binders”) for each antigen for training, so that none of our models were provided at any time with information about the top 1% strong binders. We split the weak binders into an 80% training set and a 20% test set. We trained either the variational Bayesian neural network (vBNN) used in our previous RESP1.0 pipeline^12^ or the approximate Gaussian process implemented by us in the xGPR library in more recent work^28^.

Full results appear in the Supplementary Info under table S1 and Figure S1. In every case, regardless of whether R^2^, mean absolute error (MAE) or root mean squared error (RMSE) is used as a metric, xGPR outperformed the vBNN by large margins. Across all 13 antigens, xGPR on average achieves an R2=0.39 higher than that of the vBNN (95% CI, +/− 0.15). Further, xGPR exhibited improved uncertainty calibration as demonstrated by a significant reduction in AUCE. On average across all 13 antigens the AUCE for xGPR is 0.23 lower than the AUCE for the vBNN (95% CI +/− 0.07) (for AUCE figures and confidence intervals see Supplementary Info table S1). These results demonstrate the superiority of the xGPR uncertainty-aware model we use in this paper in preference to the vBNN used in RESP1.

Next, we used the trained xGPR model to conduct RESP searches for CDRH3 sequences likely to tightly bind all of the target antigens in a group. For each antigen group, we ran 20 RESP searches using a different random seed and randomly selected starting sequence from the training set for each (see Materials and Methods, section “Experiments on synthetic data from the Absolut! Database” for details). We retained only candidates predicted by the model to be tighter binders than any sequence present in the training set. Crucially, we filtered the candidates generated by the RESP search to remove those for which the model has low confidence (see Materials and Methods, section “Experiments on synthetic data from the Absolut! Database” for details). Even after applying these filters, the xGPR-powered RESP searches generated > 75 candidates for all protein targets and on average generated 484 candidates per target group.

We next compared the RESP approach with two generative AI approaches currently popular in the literature. First, we fine-tuned the 640 million parameter Evodiff-Seq model from Alamdari et al.^31^ on either the top 20% of sequences from the training set (the 80th percentile) or the top 10% of sequences from the training set (the 90th percentile) (see Materials and Methods, section “Experiments on synthetic data from the Absolut! Database” for details). The fine-tuned Evodiff-80 and Evodiff-90 models were used to generate 1,000 candidates each per antigen group. Finally, we used the Protein-MPNN model of Dauparas et al.^24^ with the starting antibody-antigen complex from the PDB to generate 1,000 candidates for each antigen in each group, modifying CDRH3 only and masking the rest of the sequence.

All candidates retrieved from the RESP search and generative AI procedures were docked and scored using the Absolut! software toolkit. The results appear in Figure 2 and in Supplementary Table S2. It is immediately clear that RESP search outperforms fine-tuned generative AI models and Protein-MPNN by large margins; the best success rate for a generative AI approach is < 1.5%, and Protein-MPNN always achieves a 0% success rate. This is consistent with the highly variable but frequently low success rates for generative AI techniques in the literature. Bennett et al.^20^, for example, used the RFDiffusion pipeline (which relies on Protein-MPNN to generate candidate sequences) to generate binders to 6 different epitopes on 5 different antigens but frequently achieved very low success rates. They tested 18,000 candidates against RSV, for example, and their best binder exhibited an affinity of 1.4 micromolar, while for SARS-Cov2 out of 9,000 candidates generated their best binder exhibited an affinity of 5.5 micromolar. Their results are consistent with what we observe here: generative AI sometimes succeeds but with very low efficiency. RESP by contrast always achieved a success rate > 84%. Given these promising results for multi-target optimization on synthetic data, we next used our updated pipeline for a real-world proof of concept with variants of the RBD of the SARS-Cov2 spike protein as targets.

**Figure 2.**
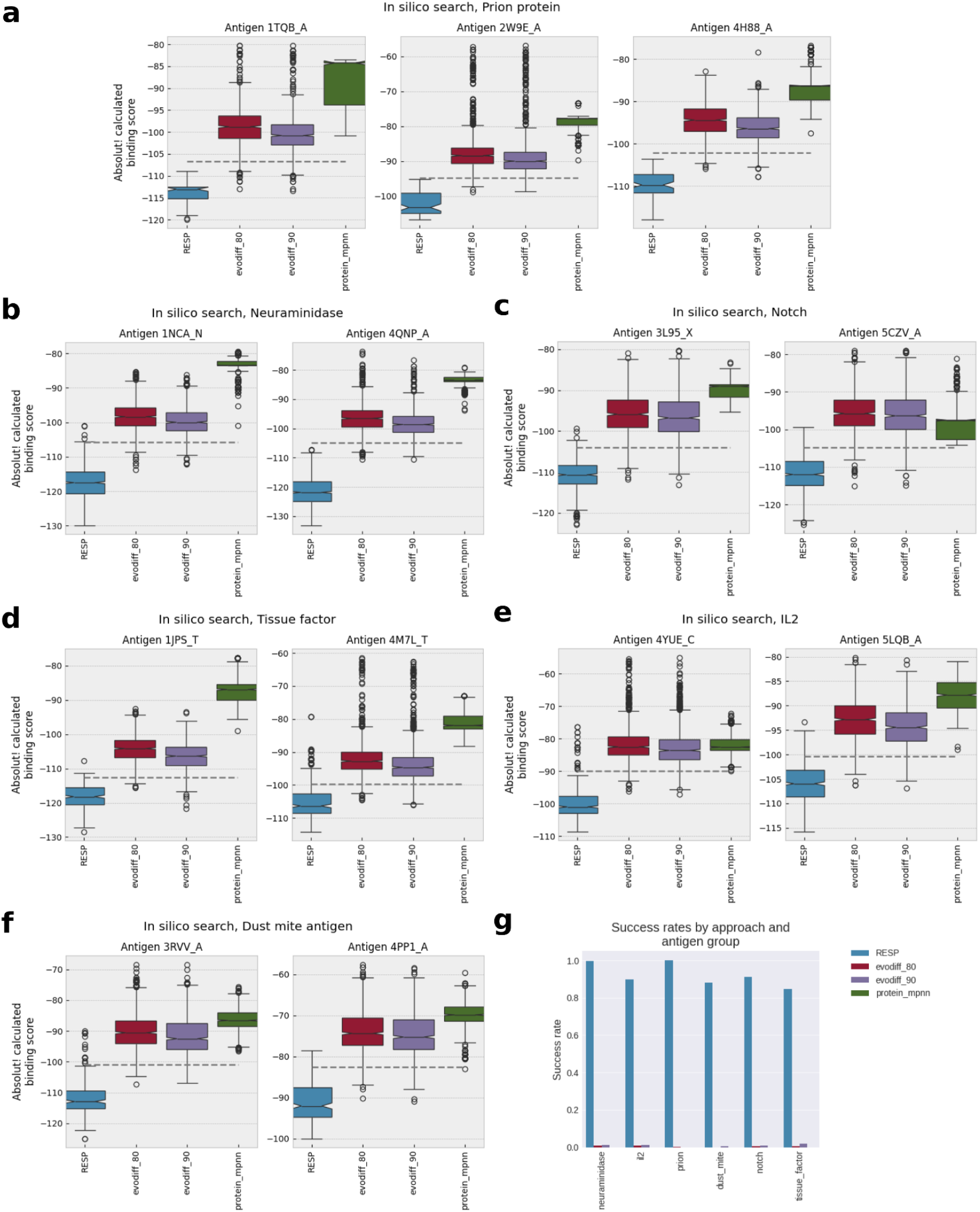
Affinity of candidate sequences for each antigen and candidate generation method in each synthetic data target group as quantified by the Absolut! docking algorithm. The lower the score, the tighter the binder. The dashed line indicates the affinity of the tightest binder in the training set. 1a. Results for the prion protein. 1b. Results for neuraminidase 1c. Results for Notch protein. 1d. Results for tissue factor protein. 1e. Results for IL-2. 1f. Results for dust mite antigen. 1g. Overall success rates, where success is defined as the fraction of candidates generated that bind more tightly to all antigens in a target group than any sequence in the training set.

### Generation and Screening of the scFv Heavy Chain Library for Improved Binding

The heavy chain of the starting clone (Delta-6 scFv, see Supplementary Figures S2 & S6 for its sequence) was mutated randomly by error-prone PCR using a previously described method^32^, creating an average of 2 mutations per heavy chain. It is of course possible to modify both the heavy and light chain using the procedure we outline here. In our previous experience, however, modifying the heavy chain only can be sufficient to achieve high affinity. In our previous work^12^, for example, we identified an scFv with a 5.3 pM K_d_ by modifying the heavy chain of an existing antibody drug against PD-L1. Moreover, modifying the heavy chain only facilitates deep sequencing, since the length of the region that must be sequenced is thereby reduced by roughly 50%. We therefore modify the heavy chain only in these experiments, although the light chain can also be re-engineered if desired. The starting Delta-6 clone, which was derived from a hit from a naive yeast display scFv library^33^ in our lab, was selected because it bound the RBD of the Delta variant (which was prevalent during the earlier stage of this project) with a K_D_ of 1.3 nM (Figure 5C). We mutated the heavy chain region of the starting clone using error-prone PCR to generate a library with an estimated diversity of at least 10^7^ members.

This Delta-6 scFv library was then screened against 10 different RBDs (SARS COV-2 WT, Alpha, Beta, Gamma, Delta, Lambda, Kappa, B.1.1.529, BA.2, and BA.5,) incubated at various concentrations with the yeast library and sorted by FACS into different bins (see supplementary figure S5 for examples of FACS plots). If the Delta-6 scFv already had lower nM affinity against the RBD (WT/Alpha/Delta/Lambda), then the concentration of RBD used in the sort was approximately equal to the K_D_ value and the hits sorted into either a “WT-level” bin (about the same affinity as the WT Delta-6 scFv) or a stronger-affinity bin. The hits corresponding to the WT-level were gathered to scrutinize the heavy-chain positions that appeared to be invariable, essential for preserving a WT-level (low nM) K_D_ with the specific RBD (i.e., positions to avoid mutating in a focused library due to high conservation). For RBDs with poorer/no apparent affinity to Delta-6, the hits were binned into either a “better than WT Delta-6” group or “significantly better than WT Delta-6” group using 2 different RBD concentrations. Also, an additional 200,000 random library clones were collected to provide information about pre-gating sequence frequencies for modeling.

We found that a single sort was sufficient for RBDs with low nM K_D_ values to Delta-6 (WT/Alpha/Delta/Lambda variants) to effectively enrich for stronger binders but 2 sorts for the rest of the RBDs were necessary. After 1-2 sorts, library hits were grown to high density then harvested for deep sequencing.

### Benchmarking uncertainty-aware models predicting sequence enrichment

We first calculated the relative abundance of each sequence in each binding bin compared to its abundance in the original library (see Materials and Methods for details). We refer to this quantity as *enrichment* and use it as a proxy for binding affinity. The FACS sorting data measures the abundance of a sequence in the unselected library and also in a binding bin for each antigen. A sequence that is more common in the binding bin for a given antigen than in the unselected library is likely to bind to that antigen, whereas a sequence that is more abundant in the original library is less likely to be a binder. The antigens were experimentally tested in two separate groups as described above. Group A contained Lambda, WT, Delta and Alpha, which underwent a single sort. Group B comprised Beta, B.1.1.529, BA5, BA2, Gamma and Kappa, with two sorts applied. Landrum et al.^34^ recently demonstrated that combining measured *Ki* and IC50 values from different sources can introduce substantial noise into datasets for ML, sometimes completely obscuring the signal of interest. In this instance, the different number of sorts between the groups could be a confounding factor. To avoid any impact on the model’s reliability, we modeled each group separately.

Next, we trained a supervised learning model to predict enrichment, which as before was used as an indirect proxy for binding affinity. RESP2 provides three models for estimating uncertainty. These include the variational Bayesian neural network implemented in the original RESP^12^ and xGPR^28^, a fast random Fourier features-approximated Gaussian process (RFFAGP) with sequence kernels that could provide performance competitive with deep learning for many sequence property prediction tasks, while providing uncertainty estimates significantly more well-calibrated than popular uncertainty estimation techniques for deep learning models. In the Absolut! synthetic data experiments described above, we demonstrated the superiority of xGPR to the vBNN used in the original RESP. Gaussian processes require suitable kernels selected by the user for a given problem^35^. While appropriate xGPR kernels are available for proteins and small molecules, selecting an optimal kernel for some datasets may be non-trivia. In contrast, deep learning is much more flexible and may therefore be easier to use for new datasets. Consequently, we additionally provide in RESP a new model called ByteNet-SNGP that combines deep neural network with RFFGP by using the RFFAGP as the last layer of the deep learning framework.

As demonstrated by Li et al.^36^, when used in conjunction with spectral normalization on the neural network weights, known as spectral-normalized Gaussian process (SNGP), this model can achieve well-calibrated uncertainty estimation and is compatible with any deep learning architecture. The deep learning model provides great flexibility to learn a representation of the inputs which is well-suited for the RFFAGP layer to achieve optimal performance. Therefore, this option unlike a Gaussian process does not require kernel selection and can provide greater flexibility. Ko et al. explored the use of SNGP-quantified uncertainty for protein-protein interaction prediction in recent work^37^; the use of the SNGP for protein and antibody engineering has not otherwise been studied previously.

We considered two possible deep learning architectures in combination with the SNGP. The first was the well-known transformer architecture^38^. The second was the ByteNet convolutional neural network (CNN)^39^. In recent work, Yang et al.^40^ demonstrated that a ByteNet which “learns” to reconstruct its input provides performance highly competitive with language models like ESM-2, suggesting that the ByteNet CNN architecture could serve as an effective general-purpose model for protein sequence data. The core structure of ByteNet (Figure 1B; for more details, see Materials and Methods) employs progressive convolutions with increasing dilation to capture long-range dependencies between all parts of a sequence. We adapted this architecture to process two paired sequences – an antibody and an antigen – as inputs, which provides a new flexible uncertain-aware model for antibody engineering.

The models we use here are trained to predict the enrichment (a proxy for binding) of a specific input antibody sequence for a specific input antigen sequence. Each model therefore accepts as input an encoded antibody and an encoded antigen, so that the model “learns” how mutations in both the antibody and antigen sequence affect binding. For xGPR and the vBNN, the antibody and antigen representations are concatenated. The ByteNet SNGP model processes the antibody and antigen representations through two separate ByteNet blocks then concatenates the output of those blocks as illustrated in Figure 1. See Materials and Methods for more details.

We evaluated these models alongside a range of encoding schemes. The models under consideration included 1) linear regression, 2) variational Bayesian neural network, similar to the one in RESP^12^, 3) xGPR with an RBF kernel but with different length scales for the antigen and antibody sequences, approximated using 32,768 random Fourier features, 4) xGPR with the FastConv1d convolution kernel, a kernel developed for modeling protein sequences^28^, 5) ByteNet deep learning architecture^39^ with 10 layers, either non-uncertainty aware (the standard architecture) or uncertainty aware (with an RFFAGP last layer, i.e. an SNGP^36^). Additionally, we explored a four layer transformer (i.e. a multi-head self-attention based architecture with separate modules for the antigen and antibody) but initial experiments (data not shown) indicated challenges in achieving validation set performance comparable to ByteNet. Consequently, our subsequent experiments focused on the ByteNet architecture.

In our investigation of encoding schemes, we considered 1) one-hot encoding, 2) PFASUM encoding using a common substitution matrix derived from Pfam seed multiple sequence alignments (MSAs), which is similar to BLOSUM but tailored to the known sequence space for improved homology search results and MSA quality^41^ and 3) encoding using learned representations combining the AbLang representation^42^ with the ESM1v representation^43^ for the antigen. The use of embeddings for the antibody greatly increased the memory and time cost of model training; for instance, each of 104 antibody positions is represented by a 768- and 1280-length vector in AbLang and ESM1v, respectively. There are many other embeddings we could use to represent antibody and antigen sequences, such as AbLang^42^, IGLM^44^, AntiBerty^45^, Antiberta^46^, ProGen-OAS^47^ and BALM^48^. It is beyond the scope of this paper to consider all. We use AbLang and ESM due to their frequent applications for this task. We used the ESM-1 embedding rather than ESM-2^49^ since it is not only significantly faster but also frequently provides similar performance. Notin et al., for example, found that across over 200 benchmark datasets, the average Spearman’s-r for correlation between zero-shot predicted fitness and actual fitness for ESM-1b is statistically equivalent to the performance achieved by the much larger ESM-2 15-billion parameter model^50^.

We randomly split the raw data into 80% train, 20% test. For linear regression and the Gaussian process, we tuned kernel hyperparameters by maximizing the marginal likelihood on the training data, using the Python xGPR package^28^. For all other model architectures, we tuned the hyperparameters of the model by evaluating performance on a held-out subset of the training data. Finally, we assessed performance for all models on the held-out test data. Results are summarized in Figure 3A; details and the error bars on each result appear in the Supplementary Info Table S3.

**Figure 3.**
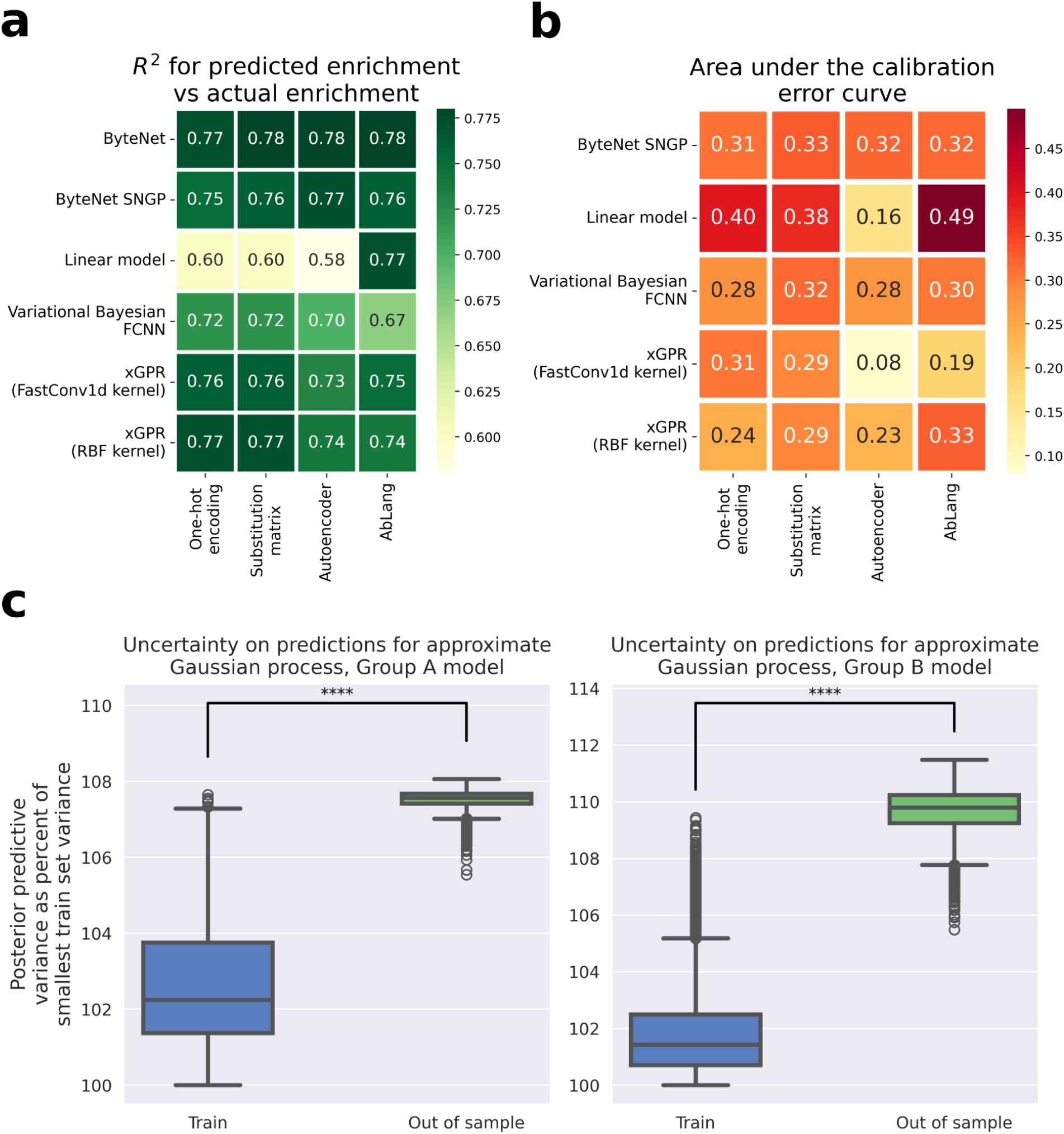
3A. Test set performance for selected model architectures and data encoding types (see Supplementary Info Table S3 for error bars on the R^2 values and for mean absolute error (MAE) and root mean squared error (RMSE) scores). 3B. Area under the calibration error curve (AUCE) for selected model architectures and data encoding types. 3C. Uncertainty on decoys (sequences known to lie outside the region of sequence space covered by the training set). We generate 10,000 “out-of-sample” mutants by randomly mutating the starting heavy chain sequence in 20 locations. We then calculate the posterior predictive variance on these sequences vs the variance on the training and validation sets using the final trained random features approximated Gaussian process models for group A and group B antigens. In both cases, the model’s uncertainty for decoys is significantly greater than that for non-decoys. In both cases, using the two-sided Mann-Whitney U test as implemented in Python’s Scipy library version 1.5.4, the calculated p-value for training vs out of sample and for validation vs out of sample is 0.0. The following conventions apply for each boxplot. The upper and lower bounds of the box are the 25th and 75th percentile of the data, and the whiskers are drawn at 1.5x the interquartile range (the distance from the 25th percentile to the 75th percentile). The center is drawn at the median of the data. The circles represent “outlier” points which lie outside 1.5x the interquartile range. Four asterisks indicate the p-value is < 0.00001. Source data are provided as a source data file.

When performance is quantified using R^2^ or root-mean squared error (RMSE). xGPR achieved performance generally similar to the more complex ByteNet-SNGP model, and the vBNN and linear regression are worse than either xGPR or ByteNet. The performance of the ByteNet-SNGP was slightly improved by using embeddings for the antibody as input, at the expense of a dramatic increase in computational expense. xGPR had marginally decreased performance when embeddings are employed, likely because significantly more random Fourier features are needed to work well with an extremely high-dimensional input. Surprisingly yet consistent with our past findings^12^, one-hot encoding achieved comparable performance. If performance is quantified using mean absolute error (MAE), which is less sensitive to poor predictions for outliers, ByteNet is superior.

To evaluate the quality of uncertainty estimation, we calculated the area under the calibration error curve (AUCE)^51^ for the ByteNet-SNGP, the xGPR and the vBNN using one-hot encoding. AUCE varies from 0 to 1, where 0 indicates perfectly calibrated uncertainty and 1 represents the poorest calibration possible. While all three models achieved comparable results, the xGPR appeared to be better-calibrated (Figure 3B and Supplementary Info, Table S4).

This comparative analysis indicates that xGPR and ByteNet-SNGP showed better performance and calibration of uncertainty estimates than the variational Bayesian fully-connected NN used in the original RESP pipeline. xGPR was efficient for this task, fitting the data in about a minute or less, a stark contrast to ByteNet-SNGP which required several hours. Furthermore, hyperparameter tuning for xGPR is completed in time frames ranging from < 30 minutes to up to two hours, depending on the expense and rigor of the hyperparameter tuning procedure selected. ByteNet-SNGP by contrast took a week of experiments.

Based on these results, we one-hot encoded the antibody. We then encoded the antigen using an ESM1v embedding averaged over the sequence (i.e. the embeddings for each amino acid are averaged to yield a single vector representation). We then fitted the data using xGPR as above.

To further confirm that uncertainty assigned by xGPR could detect out of distribution data, we generated a dataset of 10,000 mutants by introducing 20 random mutations into the wild-type sequence. Given that the average training and test set sequence had only 3.7 mutations, we expected the model to be significantly more uncertain about these new sequences, which it was (Figure 3C, p = 0.0, Mann-Whitney U test). These results appear in Figure 3.

### Identifying important positions for *in silico* directed evolution

To identify positions most likely to improve affinity, we mutated each position *in silico* to each possible amino acid (except cysteine) and assigned scores against all antigens using the model. The smallest score for any antigen was used as the score for that mutation, and the largest score for any mutation at each position was used as the score for that position. This approach identifies positions where there is at least one possible mutation that yields a large score increase against *all* antigens (rather than only one or a few of them).

The choice of the number of positions depends on available experimental resources; the larger the number of candidates that can be tested, the larger the search space that can be adopted. In our case, we sought to limit the number of candidates to < 30 given our available experimental resources. To limit the search space for *in silico* directed evolution, we picked the top six highest scoring positions. This procedure yielded positions 26, 27, 29, 32, 73 and 103 as the best positions for mutagenesis. 26, 27, 29 and 32 are all in CDR1, while 103 is in CDR3. For simplicity, both here and elsewhere throughout this paper, we designate positions by numbering sequentially from the first position in the mutated heavy chain, and define CDRs and frameworks using the IMGT numbering scheme and CDR definitions^52^. Position 73 is the only one outside a CDR. To assess the risk associated with modification of this position, we evaluated the human-likeness of the wild-type sequence using AntPack (v0.2.7). We found that depending on which mutation was selected, this framework position could be mutated with a decrease in human-likeness score of < 3%, and that certain mutations at this position are therefore likely to have negligible impact since the sequence and framework with this modification is still rated as highly human by the model.

When using an automated procedure such as the one we describe here, it is important to enable human oversight so that practitioners can double-check the decisions made by the model. To explore the raw data in greater detail and to provide a “sanity check” on important position identification, we also fitted a mixture of categorical distributions to sequences with enrichment values of 3x or above in two or more bins, i.e. sequences for which the probability of the sequence in a binding bin is at least 20x greater than that in the naive bin for two or more antigens. This simple model calculates the probability of any sequence of length L as:

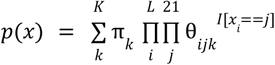

Where the sum is across *K* clusters π*_k_* is the mixture weight for each cluster, θ*_ijk_* is the probability of amino acid *j* (21 amino acids including gaps) at position *i* for cluster *k*, and I is the indicator function. Each cluster treats positions as independent, but since probabilities are conditional on cluster membership, the full model captures relationships between positions and the joint distribution. The model is fully interpretable and provides a simple yet powerful tool for dataset exploration. This model’s simplicity makes it very easy to fit quickly; Parkinson et al. for example fit it to 70 million light chains and 60 million heavy chains^29^. Despite its simplicity, this type of model has not previously been used for protein engineering to our knowledge. We selected the number of clusters by finding the number which minimizes the Bayes information criterion (BIC) (for more details on the fitting procedure, see Materials and Methods). This procedure yielded 9 clusters.

The probability of each possible mutation in each cluster is illustrated in Figure 4, with the CDR regions shaded. Every single cluster features either the F27L or F27I mutations with probability 0.98 or above (for seven of the clusters, > 0.99), suggesting the F27 mutations are key for improved affinity. Two clusters feature a G26E mutation with probability 1 and a third exhibits G26E with moderate probability (> 0.3). Positions 103, 28 and 73 have a low-moderate probability of mutation in some clusters (for example, position 73 is mutated with about 25% probability in cluster 1).

**Figure 4.**
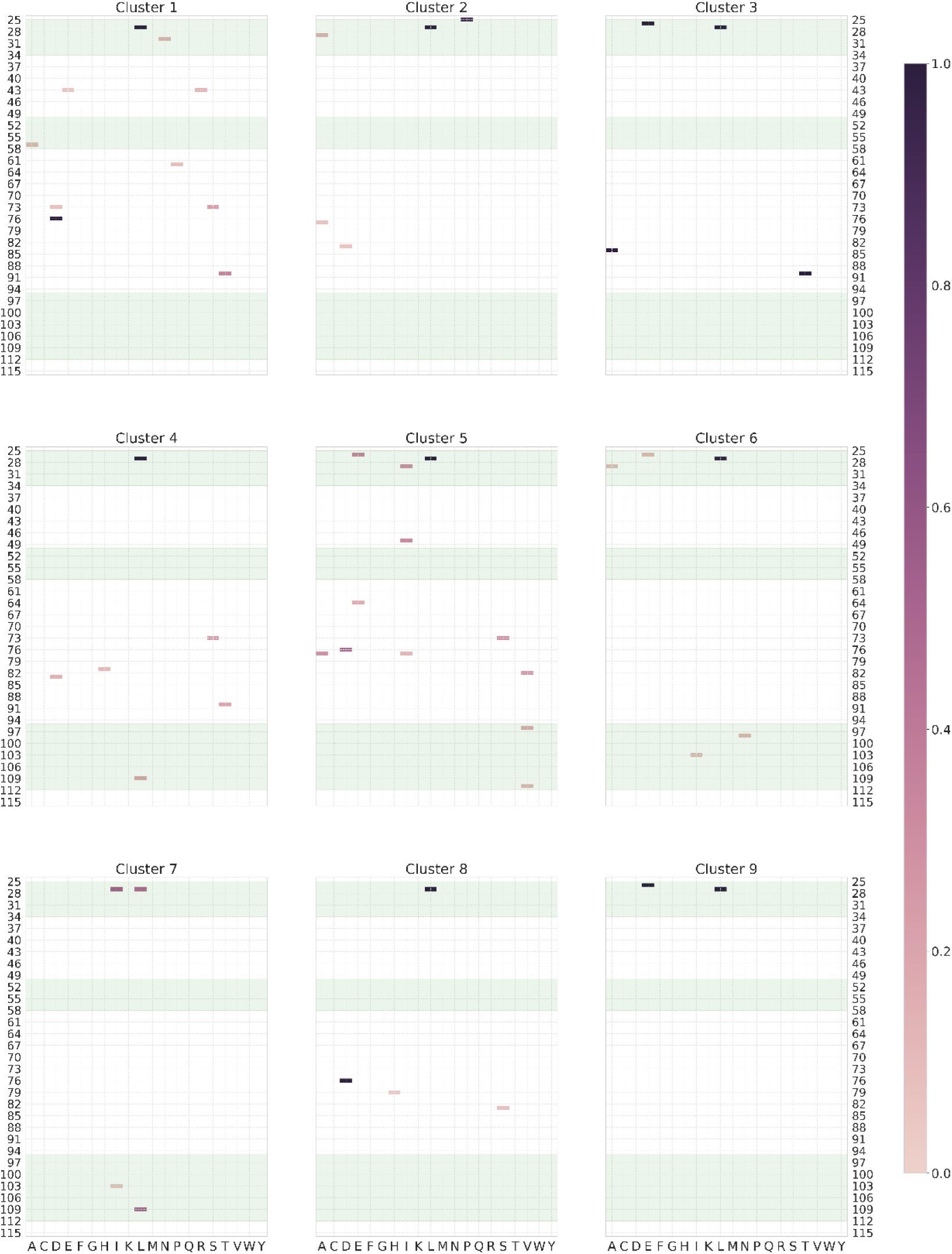
The probability of each mutation in each cluster for a mixture of categorical distributions fitted to highly enriched sequences. Positions are numbered sequentially from the first position in the modified heavy chain sequence.

There are two interesting discrepancies between the mixture model-preferred positions and those selected by the automated procedure described above. Position 31 is not identified as a key position by the mixture model, indicating it is not mutated in the most highly enriched mutants but only rather in more moderately enriched mutants. A more interesting discrepancy involves position 76. The mixture model identifies this as a promising position for mutation, with a mutation probability > 99% in two clusters. The affinity model also predicts large score increases > 1 for most antigens if this position is mutated, but predicts smaller score increases (< 0.8) for antigens B.1.1.529, ba5 and ba2, and moderate score increases for beta and gamma.

Interestingly, the affinity model further predicts that mutating 76 *in combination* with the F27L mutation will give a much larger increase in score for B.1.1.529, BA5 and BA2 than either mutation alone. The two mutations in combination result in a predicted score increase > 4 for all antigens. Both models therefore concur that mutating position 76 is likely beneficial in the presence of certain other specific mutations but is not likely sufficient (note there are no mixture model clusters in which 76 is mutated without a corresponding mutation at position 27). Modifying 76 would however incur some additional risk since the selected positions already include another framework position at 73. In light of this consideration we used the set of positions selected by the automated procedure we describe above. The workflow we have outlined here illustrates how practitioners can model the distribution of mutations in tight-binding sequences in a fully interpretable way and choose to modify the positions selected through a simple automated procedure as appropriate.

### *In silico* directed evolution to select the most promising candidates

In RESP2, after training a model to predict affinity, we employ it to conduct *in silico* directed evolution (ISDE) using a modified simulated annealing algorithm. During each iteration, the current candidate is mutated at a randomly selected site and accepted with probability:

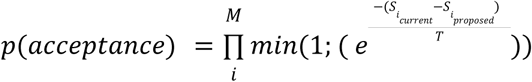

Where *T* is the temperature and *S*_*current*_ and *S*_*proposed*_ are the current candidate score and the score of the proposed sequence respectively, *i*… *M* runs over the *M* antigens in the panel. The scores here are the predicted affinities / enrichments from the affinity model. We introduce a novel modification to typical simulated annealing, however, by rejecting any candidates for which model uncertainty on its predicted score is high. This key modification ensures the directed evolution process does not wander into regions of sequence space where model predictions are unreliable. This strategy selects sequences that exhibit improved binding to at least one antigen on each round.

To ensure good developability (low risk of immunogenicity, good solubility, stability etc.), we filter the sequences generated by ISDE for solubility (using the CamSol model v2.2, available at https://www-cohsoftware.ch.cam.ac.uk//index.php^53^) and for humanness (using the SAM model of Parkinson et al.^29^).

The SAM model exhibited superior performance on a test set of 400 million sequences for distinguishing human heavy and light chains from those of other species^29^. Importantly, since it is fully interpretable, it can suggest modifications that would improve humanness for a tight binder with initially low scores. These modifications are then evaluated by the affinity model to ensure they do not compromise binding, facilitating a complete *in silico* humanization process.

In this study, we conducted five ISDE experiments and harvested 116 sequences, which still far exceeded our testing capacity. Given the available resources, our objective was to select < 30 sequences for testing. We removed any with low humanness scores or high relative uncertainty in model-assigned score (for details on the selection criteria, see Materials and Methods), then selected the 29 remaining sequences with the highest model-assigned score.

### Validation of the predicted tight binding antibodies

We transformed all 29 selected candidates into yeast to generate thousands of transformants. We also transformed the WT Delta-6 scFv as a control for the flow cytometry experiments.The WT Delta-6 scFv or the 29-member mutant library were grown and induced and incubated with the 10 different RBDs. For the RBDs with measurable K_D_ values to Delta-6, the K_D_ was used as the labeling concentration, while for RBDs with no measurable K_D_ to Delta-6 the concentration was set at 321 nM, the highest concentration used in the K_D_ measurements.

The flow cytometry plots showed that the 29-member library exhibited a significantly higher binding signal than the WT scFv control for 10/10 RBDs (see Figure 5A for example FACS plots, supplementary figure S7 for all the plots). The percentage of library members with higher intensity than the WT ranged from 67.4% for the Gamma RBD to 90.6% for the Alpha RBD (see Figure 5B for full listing of % library members more intense than WT Delta-6). Importantly, there was minimal non-specific binding to the secondary detection reagent streptavidin-PE with 2.2% events positive for Delta-6 compared to 2.9% for the 29-member library (fig. S7). This indicates that the improved binding of the library is due to specific interaction with the RBD rather than non-specific (like hydrophobic or charge-charge) effects.

**Figure 5.**
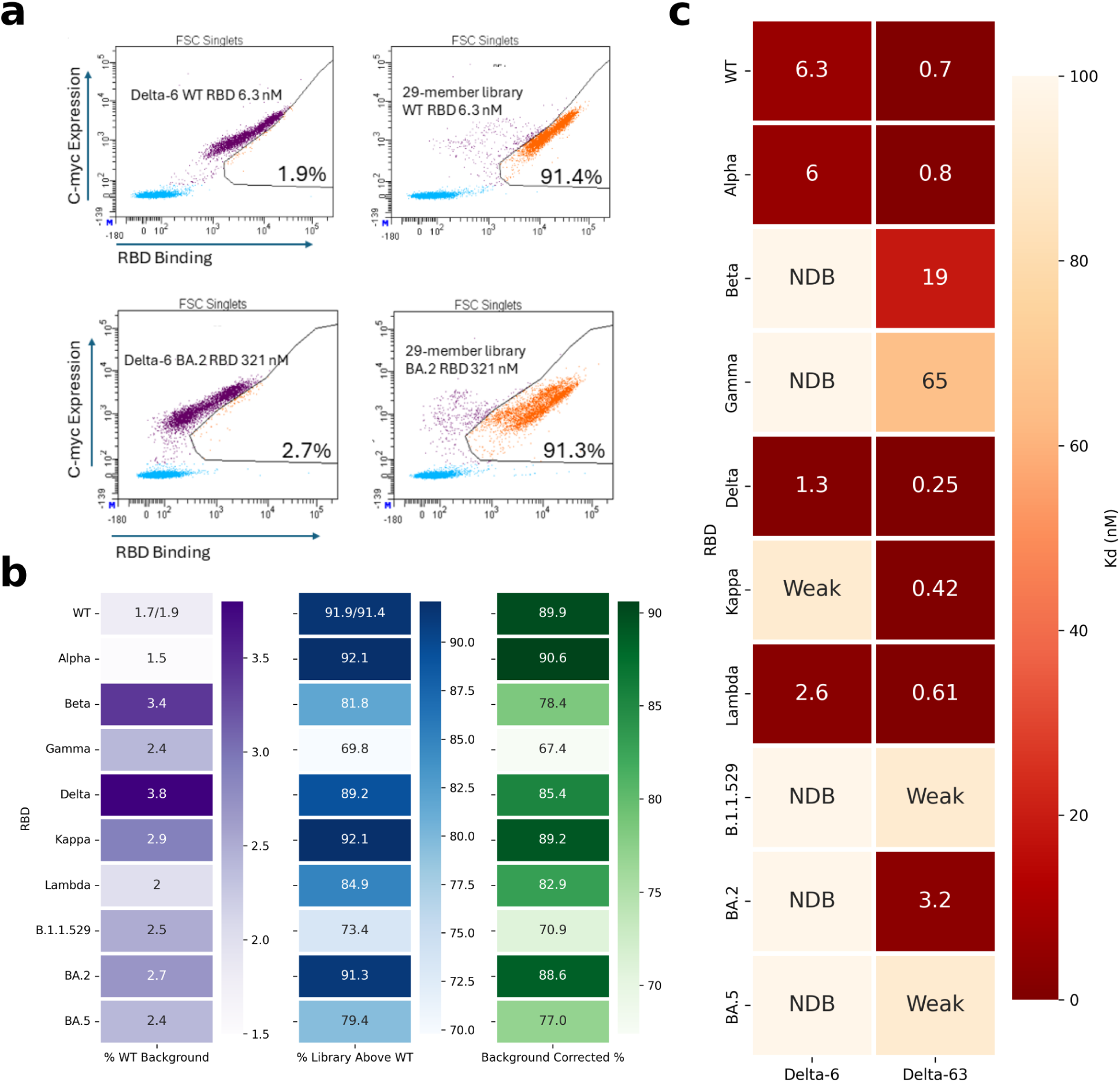
5a. Examples of flow cytometry analysis of WT Delta-6 scFv vs. 29-member library using the RBDs of WT SARS COV-2 (top panels) and the BA.2 variant (bottom panels). The % values represent % of expressing library members above the WT Delta-6 main population. As a control, the % of WT Delta-6 cells appearing above the WT main group is included. The y-axis of each plot represents scFv expression (c-myc tag) while the x-axis the binding intensity to the RBD (PE fluorescence). 5B. The percentage of scFv-expressing 29-member library yeast cells scoring as more intense than the WT Delta-6 scFv cells for binding to each RBD, corrected for background WT cells. The WT SARS COV-2 RBD data is averaged from 2 replicates. 5C. Binding Affinity of the Delta-6 or Delta-63 scFv on the yeast surface towards the 10 RBDs used in the Delta-6 library screen. NDB is no detectable binding, while “weak” means the K_D_ was too weak to determine a numerical value but binding was observed. A single replicate was performed for each K_D_ value.

To avoid the expense of measuring K_d_ for all 29 candidates, we picked one of the 29 clones, named Delta-63, at random and measured its K_D_ values individually against each of the 10 RBDs, comparing them to those of Delta-6 (Figure 5C). Remarkably, Delta-63 demonstrated tighter K_D_ values than Delta-6 for all RBDs tested. For the RBDs of B.1.1.529 and BA.5, we could not determine precise K_D_ values for Delta-63 but the binding curves clearly showed enhanced binding relative to the WT Delta-6 (see supplementary figure S8 for K_D_ plots per RBD). For 8/10 RBDs, the K_D_ values improved significantly, ranging from modest 4-5 fold increase for the Lambda and Delta variants to several orders of magnitude for the Kappa and BA.2 RBD. Specifically, the Delta-6 K_D_ values to Kappa and BA.2 RBDs were too weak to quantify but Delta-63 exhibited a marked improvement, achieving low nM to pM levels. The BA.2 variant presented a stark contrast; Delta-6’s binding was virtually undetectable, whereas Delta-63 achieved a K_D_ of 3.1 nM. Similarly, the Beta and Gamma RBDs, which initially showed negligible binding to Delta-6, displayed substantial gains of binding to Delta-63, with K_D_ values of 19 and 65 nM, respectively.

We further tested if Delta-63 could inhibit the RBD-ACE2 interaction. When 3.2 nM of the WT SARS COV-2 RBD was added to yeast cells expressing Delta-63, a robust binding signal was observed. This signal was reduced more than 3 fold upon the addition of a large molar excess of ACE2 (Supplementary Figure S3), suggesting that Delta-63 binds at least part of the ACE2 epitope on the RBD. This observation indicates that Delta-63 could be further optimized into a functional anti-COVID monoclonal antibody capable of inhibiting the RBD-ACE2 interaction - a characteristic of effective anti-COVID monoclonal antibodies^54,55^.

Furthermore, we measured the expression level of Delta-63 on the yeast surface, as the expression level of proteins on the yeast surface has been shown to correlate with thermal stability/soluble secretion levels^56–58^. Compared to Delta-6, alongside other COVID antibodies and an irrelevant anti-PDL1 antibody (Atezolizumab), Delta-63 had the highest expression on the surface of yeast cells (see Supplementary figure S4). Furthermore, at the highest concentrations of RBDs used to determine the K_D_ of Delta-63 and Delta-6, Delta-63 consistently showed a higher binding signal at saturation (Supplementary figure S8). This indicates higher levels of active Delta-63 scFv on the yeast surface compared to Delta-6 (again suggesting higher levels of solubility/stability of Delta-63 compared to Delta-6). This indicates Delta-63 may be a particularly stable/soluble scFv and useful for future affinity maturation campaigns against new SARS COV-2 variants.

## DISCUSSION

Here we present RESP2, a new AI pipeline for antibody engineering. We enhance the RESP pipeline with implementation and development of two new state-of-the-art uncertainty-aware models, the fast approximate Gaussian process (GP) implemented in the xGPR library and the ByteNet CNN architecture combined with a last-layer Gaussian process and spectral normalization (ByteNet-SNGP). Neither model has been used for protein engineering previously to our knowledge and both models outperform the vBNN in the RESP framework. Notably, these models provide well-calibrated uncertainty estimation. While xGPR’s uncertainty estimation is more accurate, the area under the curve of the receiver operating characteristic (AUCE) remains comparable for both models. This improved uncertainty estimation enhances the selection of promising candidates for subsequent optimization, expediting the development process.

RESP2 incorporates several uncertainty-aware models, enabling users to choose the most fitting option for their applications. The Conv1dRBF kernel in the xGPR software package works well when each datapoint is associated with only a single sequence, particularly useful when the goal is to enhance affinity against only a single target antigen, but harder to adapt to a pair of sequences when optimizing affinity against multiple antigens. Instead, as we have demonstrated here, the FastConv1d and RBF kernels are easily adapted to this task, making them viable choices. A CNN with a last-layer Gaussian process (i.e. an SNGP) is an alternative that may improve accuracy and can be selected instead if it provides substantially better performance.

Another important feature in RESP2 is the ability to develop antibodies against multiple mutated variants of an antigen, which is unique among machine learning-assisted pipelines for antibody development. The modified *in silico* directed evolution process considers model predictions for a multitude of antigens prioritizing sequences likely to have improved binding across all targets. We also introduce a simple yet flexible and fully interpretable mixture model for exploring the raw data. The mixture model identifies the positions that have the highest mutation probability in tight binders and thereby serves as a valuable “sanity check” for fully automated position selection.

Finally, we incorporate models for predicting developability. We make use of a fully human-interpretable generative model for human antibody sequences implemented in the AntPack package^29^, which can distinguish human from nonhuman sequences with better accuracy than any other comparator model currently available. As demonstrated in this paper, we can make use of humanness scores provided by AntPack at two stages in the pipeline. First, we can select positions for *in silico* mutagenesis where a mutation would be unlikely to impact humanness. This is generally true for CDRs, of course, but we can use AntPack to identify mutation-tolerant positions in the framework as well. Second, we can filter candidates generated by the pipeline based on their humanness scores, removing any which are suboptimal. We have included solubility prediction using the CamSol model^53^, and the pipeline can readily accommodate additional computational filters as desired. For example, it is easy and straightforward to incorporate predictions for MHC class II binding, which can further lower the risk of immunogenicity by removing mutations that increase risk.

To demonstrate the power of RESP2, we experimentally assess our pipeline both *in silico* and on real data. We first demonstrate that unlike generative AI techniques, RESP2 consistently achieves very high success rates on synthetic data against six groups of antigens. Next, we use the COVID-19 RBD as a test case and successfully evolve a weaker-binding antibody to one exhibiting low nanomolar to subnanomolar affinity against multiple targets after constructing only a single random library. Concurrently, we are able to optimize developability properties. The capability to simultaneously improve binding against multiple antigens and developability is not currently practicable for any purely experimental technique.

## MATERIALS AND METHODS

### Media Recipes

PBS used for the yeast experiments was 140 mM NaCl, 2.7 mM KCl, 10 mM Na_2_HPO_4_, 1.8 mM KH_2_PO_4_, pH 7.4. SDCAA plates used to titrate yeasts are made with 5 g/L Bacto Casamino Acids (Thermo Fisher 223050), 20 g/L D-(+)-Glucose (Sigma G8270), 1.7 g/L yeast nitrogen base w/o amino acids or ammonium sulfate (BD Difco 233520), 5.3 g/L ammonium sulfate (Fisher A702-500), 10.19 g/L Na_2_HPO_4_-7H20 (VWR 0348-1KG), 8.56 g/L NaH_2_PO_4_-H20 (VWR 0823-1KG, 182 g/L D-sorbitol (Sigma S1876-500G), 16.67 g/L BD Bacto Agar (214010), and 100 μg/mL ampicillin. SDCAA citrate media (same recipe as for SDCAA plates except no sorbitol, agar, and 7.4 g/L Citric Acid Monohydrate (Macron 0627-12) + 11.85 g/L Sodium Citrate Dihydrate (MP Biomedicals) instead of the sodium phosphate reagents) and no ampicillin. SGCAA media for induction of surface expression is the same as the SDCAA plate formula but with no agar/sorbitol and 20 g/L D-(+)-Galactose (Millipore Sigma G5388) instead of glucose. Generally, 100 μg/mL ampicillin (final concentration) was freshly added to the SGCAA media. For transformation, EBY100 cells were grown in 2xYPAD media before heat shock (2% Bacto Yeast Extract (BD 212750), 4% Bacto Peptone (BD 211677), 160 mg/L Adenine hemisulfate salt (Sigma A3159-5G), 4% D-(+)-Glucose (Sigma G8270-10kg)).

### scFv K_D_ determination on the yeast surface

The dissociation constant (K_D_) was determined basically as described^32^. Specifically, 10^5^ yeasts were added per Eppendorf tube and serially diluted RBD (Acro Biosystems #SPD-C82E9, SPD-C82E5, SPD-C82E7, SPD-C82E6, SPD-C82Ec, SPD-C82Ed, SPD-C82Eh, SPD-C82E4, SPD-C82Eq, and SPD-C82Ew) were added per tube (diluent was 1%-BSA (Biotium #22013) in PBS. Once RBD was added, tubes were rotated at RT overnight. The following morning, 1:100 dilution of anti-c-myc mAb (Cell Signaling Technology 2276s) was added and tubes rotated for another hour at RT. Afterwards, tubes were pelleted, yeasts washed with 1% BSA-PBS (cold), then resuspended in 100 uL PBS-BSA with 1:100 PE Streptavidin (BD 554061) and 1:100 goat anti-mouse AF647 secondary antibody (Invitrogen A21241). Cells were tapped to mix and kept on ice 20 minutes in the dark, with occasional tapping. The cells were then washed again with cold PBS-BSA and resuspended in 120 uL PBS-BSA for flow cytometry. For each data point, the PE mean fluorescent intensity (MFI) was recorded for the expressing cells. MFI was plotted along with RBD concentration to generate binding curves, fit using GraphPad Prism 10.2.1 software using the following equation: Y=Bmax*X/(Kd+X), where Bmax is the curve plateau and X each RBD concentration. The MFI without antigen was subtracted from each concentration data point to account for non-specific binding.

### ACE2 Competition Assay on the Yeast Surface

The Delta-63 scFv expressed on the yeast surface was incubated for 135 minutes at RT with either 3.2 nM WT SARS COV-2 RBD or 3.2 nM WT RBD (Acro Biosystems SPD-C82E9) + 294 nM soluble ACE2 (Sino Biological 10108-H08H) in 1% BSA-PBS (100,000 cells per tube, 1:100 dilution 2276s anti-c-myc antibody). Afterwards, cells were pelleted/washed with cold PBS-BSA and treated/analyzed as in the K_D_ determination experiments.

### Generation of the Delta-6 scFv Heavy Chain Library

The heavy chain of Delta-6 was mutated randomly by error-prone PCR using 8-oxo-dGTP (TriLink N-2034) and dPTP (TriLink N-2037) in a similar manner as previously reported^32^, where the heavy chain PCR involved either 30 cycles of error-prone PCR with 2 μM or 20 μM each nucleotide analog to create a lower or higher mutation rate (primers “D6 ePCR F/R”,, supplementary table S7). The gel-extracted lower and higher mutated DNA were PCR amplified using Q5 Hot Start DNA polymerase (NEB) (“D6 F/R” primers), gel-extracted again (Zymo Research D4002), then used for NEBuilder HiFi DNA assembly (M5520A) along with the linearized WT Delta-6 scFv in pCTCON2 (linearized by PCR with Q5 Hot Start (NEB M0493S) and “D6 linear F/R” primers, treated with DpnI (NEB R0176S), then gel-extracted (Zymo Kit)). The purified fragments were used to assemble the plasmid library via HiFi DNA assembly in 14 reactions (7 for the higher error rate and 7 for the lower error rate). The reactions were then concentrated using Zymo DNA Clean and Concentrator-5 kit (D4004), eluted in nuclease-free water, and mixed together. The concentrated plasmids were electroporated into MegaX DH10B T1R E. coli (Thermo Fisher C640003) (78.44 × 10^6^ transformants determined by serial dilution onto LB-ampicillin plates). The plasmid library was harvested after overnight growth in LB-ampicillin via ZymoPure II Plasmid MidiPrep kit (D4200), eluted in nuclease-free water. Sequencing of random clones suggested an average mutation rate of about 2 mutations per heavy chain.

EBY100 yeasts were heat-shocked with 250 μg of library plasmid using a previously described method^59^ to generate ≥ 10^7^ transformants, judged based on serial dilution onto SDCAA selection plates. The library was grown/passaged in SDCAA citrate (pH 4.5) media and frozen in aliquots at −80C in freezing media (85% ddH20, 10% glycerol, 5% DMSO).

### Screening the Delta-6 scFv Heavy Chain Library Against 10 SARS COV-2 RBDs

The naive Delta-6 library was seeded at 50 × 10^6^ live cells into SDCAA-citrate media and grown to high density at 30 C, 300 RPM. To induce surface expression of scFv, the library was diluted to OD600 = 0.5, grown back to high density in SDCAA citrate and just when the cells reached high density again (OD600 = 5) the media was switched to SGCAA media with ampicillin (100 *μ*g/mL) at a starting OD600 = 0.5, shaking at 20 C, 300 RPM for 46 hours. Cells were pelleted, washed with PBS-BSA (1% w/v BSA), then resuspended in different tubes with various concentrations of RBDs (for WT SARS COV-2 RBD = 5 nM, Alpha RBD (5 nM), Lambda RBD (2 nM), Delta RBD (1 nM), Kappa RBD (40 nM). For the rest of the RBDs, the concentration was set at 130 nM. Cells were diluted in PBS-BSA to prevent antigen depleting conditions (which depended on the RBD concentration). Yeast library-RBD mixtures were rotated overnight at RT to allow for the binding reaction to reach equilibrium, followed by addition of 1:100 mouse 9B11 anti-c-myc antibody for 1 hour at RT. Cells were pelleted and washed with 1 volume cold PBS-BSA, followed by resuspension in ice-cold PBS-BSA with 1:100 streptavidin-PE (BD 554061) and 1:100 goat anti-mouse IgG2a-AF647 and incubated with flicking on ice for 20 minutes in the dark. Afterwards, cells were pelleted and washed with 1 cold volume PBS-BSA and resuspended at 10^6^ cells per 120 *μ*L for FACS on a BD FACSAria II cell sorter. Hits from the screens were collected in 1 mL SDCAA citrate. 200,000 c-myc positive naive library members were collected as a control for deep sequencing, along with 200,000 hits from the WT SARS COV-2, Alpha, Lambda, and Delta RBD screens that had WT-level affinity to observe apparently immutable positions needed to obtain a lower nM K_D_ value. Also for WT/Alpha/Lambda/Delta, cells with higher intensity than the WT Delta-6 were collected (about the top 0.2% of all cells for each RBD). For the rest of the RBDs, since they lacked a measurable K_D_ towards WT Delta-6, only the most intense cells were collected (about the top 0.2% for each one). Hits were grown to high density at 30 C, 300 RPM and frozen/stored at −80C or used for the next round of sorting.

For all but the WT SARS COV-2, Alpha, Lamda, Delta RBDs, a second sort was necessary to get significant enrichment of hits better than WT Delta-6 scFv. Hits from the 1st round were seeded in SDCAA citrate, grown to high density and induced in SGCAA media at 20C/300 RPM in the same manner as the naive library. For Beta/Gamma/B.1.1.529/BA.2/BA.5, the cells were labeled at both 130 nM and 32 nM to either enrich binders better than WT Delta-6 (130 nM concentration) or significantly better (32 nM, a more stringent condition). For the Kappa RBD, cells were instead labeled at 40 nM (better than WT Delta-6) or 8 nM (significantly better). Also, 60,000 of the “WT-level” binders to the RBDs from the 1st round that maintained WT-level K_D_ to WT/Alpha/Delta/Lambda were collected so as to reduce their numbers to more manageable numbers for the subsequent deep sequencing step. For all hits from either the 1st or 2nd round, the total number of collected hits was less than or equal to 60,000 cells (except for the 200,000 naive c-myc positive ones). All hits were grown to high density in SDCAA citrate and the yeast plasmids harvested by yeast miniprep (Zymo Research Yeast Plasmid Miniprep II kit, D2004).

### Preparation of DNA for MiSeq PE303

The heavy chain region of plasmids isolated from the yeast miniprep for each selected group were PCR amplified with “Delta-6 NGS F/R” primers (supplemental table S7) using Q5 hotstart DNA polymerase at 20 cycles (determined optimal by qPCR), followed by concentration of the PCR products by DNA Clean and Concentrator-5 kit (Zymo Research), then gel-extraction by 1% agarose gel and Zymo Research D4002 Gel Recovery Kit. The eluted PCR products were then amplified with a second round of PCR using 2x Kapa HiFi Master Mix (KK2601) and 10 cycles of PCR with the “Ad1/2” primers listed in supplemental table S7. The PCR products were purified with AMPure XP beads (A63880), eluted with TE buffer. All PCR products were analyzed by Tapestation (both High Sensitivity D5000 and D1000 to analyze size distributions of the PCR products). All samples were diluted to 8 ng/uL and mixed together at equal volumes for MiSeq PE303. The 200,000 c-myc positive naive library members collected as a control were separately sequenced by MiSeq PE303.

### Experiments on synthetic data from the Absolut! Database

We retrieved the sequences for the 159 antigens from the Absolut! database on 11/26/24 and clustered these using the hierarchical clustering algorithm implemented in Python’s scipy library with a 50% identity cutoff based on sequence alignment and restricting clusters to contain sequences in the same family. Clusters meeting these criteria were filtered to remove sequences that were near-identical (differed by five mutations or fewer) since these might represent an insufficiently challenging task.

We obtained five pairs and one triplet which are described in more detail in Supplementary Table S5. These include two variants of influenza neuraminidase, N1 and N9; mouse and human variants of IL-2; two variants from different species of the dust mite allergen; three prion protein variants; two variants (human and mouse) of the Notch protein; and two variants of the tissue factor protein that serves as a cofactor for F.VIIa. The maximum percent identity of a sequence to any other sequence in the same target group ranged from 56% to 97.2% with an average of 78%.

For each antigen, the Absolut! Database provides 500,000 CDRH3 sequences randomly sampled from the bottom 99% of the roughly 8 million CDRH3 evaluated against that antigen and the top 1% of CDRH3 sequences evaluated against that antigen, together with the docking score assigned by the Absolut! tool for each sequence.

We used only the bottom 500,000 sequence dataset (“weak binders”) for each antigen for training, so that none of our models were provided at any time with information about the top 1% strong binders. We split the weak binders into an 80% training set and a 20% test set. We trained either the variational Bayesian neural network (vBNN) used in our previous RESP1.0 pipeline^12^ or the approximate Gaussian process implemented by us in the xGPR library in more recent work^28^.

Next, we used the trained xGPR model to conduct RESP searches for CDRH3 sequences likely to tightly bind all of the target antigens in a group. For each antigen group, we ran 20 RESP searches using a different random seed and starting sequence from the training set for each. Since the RESP search procedure by default introduces point mutations (but not insertions or deletions), we randomly selected four starting sequences each with lengths 11, 13, 15, 17 and 19 from the bottom half of the training set (i.e. training set sequences with binding affinity below 50th percentile). All starting sequences are therefore very weak binders.

When conducting the RESP search, we modified the *in silico* directed evolution algorithm of Parkinson / Hard et al.^12^ to support optimization for binding to multiple targets. In the original algorithm, we began with the starting sequence and on each iteration proposed a mutation to this sequence, then used the binding prediction model to score the candidate sequence both with and without the mutation. We then calculated the following threshold:

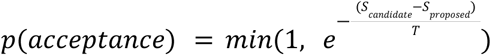

Where T is a temperature parameter initially set to 25 then multiplied by *a* < 1 on each iteration. In this way, the algorithm initially favored exploration, with a high probability that any modification would be accepted, and later favored only mutations that improve the score, since once temperature was small (e.g. < 1) the probability rapidly fell towards zero when *S*_*proposed*_ was significantly less than *S*_*candidate*_.

There are a variety of ways in which this scheme could be modified to support *M* targets. We could for example use a product:

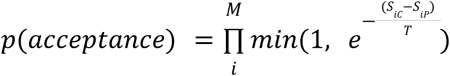

Where iC and iP denote the current candidate and proposed mutation scores for target *i*. We refer to this as the *conservative* scheme. Alternatively we could use a maximum:

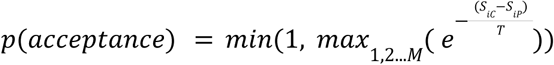

We refer to this as the *aggressive* or liberal scheme. The aggressive scheme is more likely to accept a mutation if it offers a large score improvement for any one target, even if scores against other targets are worse as a result of the mutation. The conservative scheme, by contrast, only accepts mutations with high probability if they improve or at least do not damage the score for most or all targets. See Amine et al.^60^ for a discussion of these and other alternatives for multi-objective annealing. For the synthetic data experiments, we use the “aggressive” scheme to promote finding the tightest binders possible to at least one antigen in each group.

We retained only candidates predicted by the model to be tighter binders than any sequence present in the training set. Crucially, candidates are filtered using the model’s uncertainty on its predictions, eliminating all candidates where the confidence interval on the prediction overlaps the best score in the training set and additionally eliminating the 50% of candidates with the highest uncertainty (the same procedure we used in the RESP1 paper). Even after applying these filters, the RESP searches generated > 75 candidates for all protein targets and on average generated 484 candidates per target group.

We next compared the RESP approach with two generative AI approaches currently popular in the literature. First, we fine-tuned the 640 million parameter Evodiff-Seq model from Alamdari et al.^31^ on either the top 20% of sequences from the training set (the 80th percentile) or the top 10% of sequences from each training set (the 90th percentile). The model was fine-tuned for ten epochs with a batch size of 100 and a learning rate of 1e-4 and in all cases this number of epochs was sufficient to achieve convergence (the loss on the training set had ceased to improve). The fine-tuned Evodiff-80 and Evodiff-90 models were used to generate 1,000 candidates each per antigen group. Finally, we used the Protein-MPNN model of Dauparas et al.^24^ with the starting antibody-antigen complex from the PDB to generate 1,000 candidates for each antigen in each group, modifying CDRH3 only and masking the rest of the sequence; any generated duplicates were removed prior to scoring. We used the default settings for Protein-MPNN without any modifications.

All candidates retrieved from the RESP search and generative AI procedures were docked and scored using the Absolut! software toolkit using default settings. For details on the usage of the Absolut! Software toolkit, refer to the repo at https://github.com/csi-greifflab/Absolut.

### Sequence Processing

The raw paired end reads from sequencing were checked for quality (for details of the quality filtering procedure, refer to the Supplementary Info section S1). After the quality filtering procedure, we obtained a dataset with 9,058,099 amino acid sequences or 205,520 unique amino acid sequences with an average of 3.7 mutations per sequence as compared with the wild-type.

We randomly split the raw data into 80% train, 20% test, using random seed 123, and further subdivide the training data into 80% train, 20% validation. Sequences were then encoded using the following schemes. 1) One-hot encoding for both antibody and antigen sequences. 2) One-hot encoding for the antibody sequence and ESM-1v embeddings^43^ averaged over the full sequence for the antigen. 3) AbLang embeddings^42^ for the antibody and ESM-1v embeddings for the antigen, left unaveraged with one embedding per position for the antigen. 4) The autoencoder described in Parkinson / Hard et al.^12^ for the antibody sequence and one-hot encoding for the antigen sequence. 5) A substitution matrix (the PFA-SUM matrix^41^), preprocessed so that for matrix rows *M_i_* and *M_j_* corresponding to amino acids *i* and *j*, 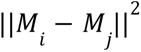 is the substitution matrix distance. Each amino acid was then represented as the row of the preprocessed matrix corresponding to that amino acid. For details on how this preprocessing was achieved, refer to Supplementary Info section S2. This last encoding is referred to as the “PFASUM matrix” in the table and is designed to work well with kernels (e.g. an RBF kernel) but is not anticipated to work well with linear regression.

We converted the frequencies of specific sequences into enrichment values that would be predicted by the model as follows. We first calculated the posterior probability of drawing each sequence from the corresponding filtered bin and the posterior probability of drawing that same sequence from the naive library using a Bernoulli likelihood with a Jeffreys prior (a Beta distribution with α = β = 1/2). The resulting posterior distribution was a beta distribution, and the posterior predictive (the probability of drawing this sequence on the next draw) was just:

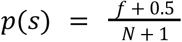

Where N is the total number of sequences in that bin and *f* is the frequency of the sequence of interest.

We then used the log of the probability in the filtered bin divided by the log of the probability in the naive bin as the *enrichment*. We also calculated the 75% highest posterior density credible interval on the posterior distribution for that sequence for both the filtered bin and the naive library. If these credible intervals overlapped, we could not be very confident that the sequence was more (or less) abundant in the filtered bin than it was in the naive library, and the datapoint was uninformative. We excluded such datapoints from our training and test sets since they did not provide useful information to the model. (We originally used a 95% credible interval rather than the 75% interval shown here, but found that this excluded over half of the training data; we relaxed the interval to 75% so that less training data would be excluded.)

The supervised learning model in the RESP pipeline^12^ was trained to predict enrichment. Positive enrichment values indicated a sequence that has some likelihood of being a strong binder, and the more positive, the more likely. Negative values, by contrast, indicated sequences unlikely to be binders. We do not need the model to be able to distinguish between sequences that are somewhat unlikely and sequences that are highly unlikely to be binders; therefore, the magnitude of the enrichment if it is less than zero does not provide useful information to the model. We therefore “clipped” enrichment so that enrichment values < 0 were set to zero.

To identify key positions, we mutated each position *in silico* to each possible amino acid (except cysteine) and assigned scores against all antigens using the model. The smallest score for any antigen was used as the score for that mutation, and the largest score for any mutation at each position was used as the score for that position. The top 6 best positions were selected. We also fitted a mixture of categorical distributions to sequences with enrichment > 3 for two or more antigens, using the EM algorithm to maximize the likelihood of the data. We used a variable number of clusters with 10 restarts with different random seeds for each number of clusters, then calculated the Bayes information criterion (BIC) for each number of clusters. The number of clusters that yielded the smallest BIC was used. The final model for each group was then fitted using 10 restarts. Any clusters with mixture weights smaller than 1e-2 at the end of fitting were purged from the model. This fitting process yielded nine clusters.

### The ByteNet-SNGP architecture and hyperparameter tuning

The ByteNet architecture uses a fully connected layer to map the input data to a length-21 embedding, so that each element of the input sequence is represented by a length-21 vector. This initial step ensures the number of parameters in the remaining layers of the model is the same irrespective of the dimensionality of the input encoding, thereby ensuring a fair comparison between different encoding or embedding types. This length-21 embedding is then fed into a series of ByteNet blocks as illustrated in Figure 1C; for this task, we used 10 ByteNet blocks. Each successive block uses dilated convolutions with increasing dilations coupled with residual connections to “learn” a representation of the input which captures both short and long-range correlations.

For this task, the model must use a paired antibody and antigen sequence as input. In our architecture, the antigen and antibody sequences were processed in parallel by two separate ByteNet modules. The output of the two modules was concatenated to form a single representation. A fully connected layer applied to each position then reduced the dimensionality of this representation to 1. In a more general case where the antibody and antigen sequences are not fixed length, this step can be replaced by max or mean pooling across all positions in the sequence; these options are available to end users utilizing the *resp_protein_toolkit* for analysis of their own data. Next, normalization is applied to the concatenated single representation. This last normalization step in initial experiments turned out to be crucial; if it is omitted, training became unstable for a wide range of hyperparameter settings. Since the antibody and antigen sequences are both fixed-length, the resulting embedding was now fixed length, and was supplied as input to a fully connected layer which yields predicted enrichment as output.

To convert this model into an uncertainty-aware architecture, we merely needed to replace the last fully connected layer with an RFFAGP as illustrated in Figure 1C. To ensure the mapping from input sequence to embedding was approximately distance-preserving, we applied spectral normalization to all weight blocks in the model^61^. This ensured that datapoints far from the training set in the original space were not mapped to locations close to the training set in embedding space. Since a GP with an RBF kernel assigns high uncertainty to datapoints distant from the training set, a distorted mapping of this kind would have led the model to incorrectly assign low uncertainty to sequences very different from the original training data.

We tuned hyperparameters for this model by varying the learning rate, learning rate schedule and weight decay and evaluating validation set performance, using one-hot encoded data as input and using the generic (non-uncertainty-aware) architecture. Following this initial tuning procedure, the same hyperparameters were used for both generic and uncertainty-aware architectures to ensure a fair comparison. Learning rate schedulers and weight decay were found not to provide a significant improvement and thus were not used. The models were trained using the Adam algorithm with a learning rate of 0.001 for 150 epochs on all datasets.

### The xGPR model and hyperparameter tuning

xGPR offers a variety of kernels for sequence data including two types of convolution kernel. The Conv1d kernels, for example, assess the similarity of any two sequences *x* and *y* using the following metric:

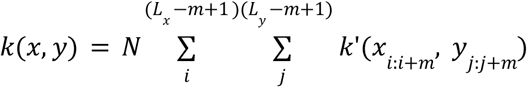

Where *x*_*i*:*i*+*m*_ is a k-mer of length *m, L* is the length of the corresponding sequence, *k*’ is a kernel function for assessing the similarity of two k-mers, and N is an optional normalization constant (to account for the number of comparisons performed). There are many possible alternatives for *k*’, but one of the simplest is an RBF kernel:

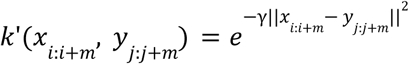

Where γ is a hyperparameter we select using an empirical Bayesian approach by maximizing the marginal likelihood of the training data. If the input data is one-hot encoded, 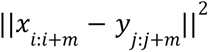 is merely twice the Hamming distance. If additionally γ is large and multiple values for *m* are selected, this is a k-mer spectrum kernel, which is closely related to a local alignment kernel that computes similarity based on local alignment; see Parkinson et al.^28^ for details. Alternatively, by encoding the input using embeddings from a deep learning model or by using substitution matrices (e.g. BLOSUM64), we obtain other measures of distance between k-mers. We can for example choose the amino acid encodings in such a way that 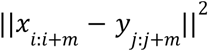 yields the BLOSUM distance between two k-mers; see the Supplementary Info section S2 for details.

A GP with the Conv1d kernels described here would naively exhibit quadratic scaling with sequence length and dataset size, but by using a fast Hadamard transform-based random features approximation, we are able to implement it in a way that scales linearly with sequence length and dataset size. See Parkinson et al.^28^ for details.

We can also use the FastConv1d kernel implemented in xGPR. This kernel conducts convolution across the sequence with random weights implemented using fast Hadamard transforms and diagonal matrix multiplications^28^, followed by ReLU activation and maxpooling. The output of this procedure is fed into an RBF kernel. This kernel constitutes an approximation to an infinitely wide three-layer Bayesian CNN populated with random weights. See Parkinson et al. for details.^28^

The FastConv1d kernel was easier to use for this particular task than the Conv1d kernels described above, since the random features generated by the FastConv1d random weight convolutions for the antigen and antibody sequences can be concatenated, while the Conv1d kernel would need to be modified to accept two paired sequences as input. We therefore used the FastConv1d kernel for convenience and concatenate the features generated for the antibody and the antigen so that the model takes both antibody and antigen variants into account.

Convolution kernels are useful when the sequences in question are of different length. In this case, however, all antibody and antigen sequences were of the same length. It is therefore even simpler to use the RBF kernel:

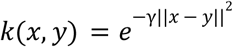

Again, if the two sequences are one-hot encoded, ||*x* − *y* ||^2^ is twice the Hamming distance. Encoding using embeddings or substitution matrices yields other measurements of distance. We can convert this kernel into a kernel for comparing antibody-antigen *pairs* by simply concatenating the antibody and antigen representations, and this is the procedure that we used in our experiments.

An obvious drawback to this kernel is that binding affinity is likely very sensitive to mutations in some regions and mostly insensitive in other regions. We can correct this problem by assigning a different γ hyperparameter to each position so that the kernel becomes:

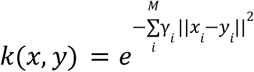

The γ hyperparameter values are found as before – by maximizing the marginal likelihood of the training data. This is implemented as a “MiniARD” kernel in the xGPR library. As for the Conv1d kernels, this kernel is implemented using a fast Hadamard transform-based random features approximation in xGPR; see Parkinson et al.^29^ for details. We refrained from assigning a separate γ hyperparameter to each position since we have found this in our experience to lead to overfitting, because the model can “learn” a small γ for a position which is never mutated in the training data but *is* mutated in the test data or in new data. A more conservative approach is to assign a different γ for each *region* of the input sequence (e.g. antibody heavy framework 1, antibody heavy CDR1, antibody heavy framework 2, antigen etc.), so that all positions in that region use the same γ-hyperparameter value.

For simplicity, in this study we assigned one γ for the antibody sequence and one γ for the antigen sequence and “learned” these by maximizing the marginal likelihood of the training data using the default approach implemented in xGPR version 0.3.2. Briefly, if the number of random features generated for each datapoint is small (<4,000), it is cheap to compute the random feature-approximated marginal likelihood exactly, without further approximations. A small number of random features is not however usually sufficient to approximate the kernel to high accuracy.

A larger number of random features can be used if the model is fitted using the preconditioned conjugate gradients (PCG) strategy implemented in xGPR, which is highly efficient and exhibits linear scaling with dataset size, but fitting the model in this way does not provide all the information needed to calculate training set marginal likelihood. If *Z* is an N x M matrix formed for N datapoints with M random features, for example, the marginal likelihood requires the calculation of the log determinant *log* |*Z*^*T*^ *Z* + λ*I*|, where λ is a model hyperparameter that regulates overfitting (analogous to the ridge parameter in ridge regression). To enable calculation of the marginal likelihood for a large number of random features, xGPR makes use of stochastic Lanczos quadrature to provide a (highly accurate) estimate of the log determinant when fitting with preconditioned conjugate gradients. See Parkinson et al.^28^ for details.

To tune hyperparameters, then, we first used the L-BFGS algorithm^62^ to maximize the marginal likelihood with 1,024 random features per datapoint; this gave a “crude” initial estimate of the region of hyperparameter space where optimal hyperparameters were likely located. Next, we “fine-tuned” the hyperparameters with the Powell algorithm^63^ using a larger number of random features per datapoint (16,384) and using the PCG-based approach described above. The xGPR documentation (https://xgpr.readthedocs.io/en/latest/) provides examples of this approach for hyperparameter tuning, which requires only a few lines of Python code.

We also fitted the data to a linear model using the “Linear” kernel in the xGPR library. We tuned the hyperparameter (the λ ridge regression parameter) by maximizing the marginal likelihood. We used an exact calculation for the marginal likelihood if the number of features was relatively small (<4,000), which was true for all encodings except the AbLang / ESM-embedding-based encoding. In this case, we set the hyperparameter to a default value of 1.

### The variational Bayesian neural network (vBNN) and hyperparameter tuning

For the vBNN, we use a three-layer fully-connected architecture as in our previous paper^12^. To supply information about both the antibody and antigen to the model, we concatenate the representations for the antibody and antigen sequences and use this concatenated representation as input. We tuned hyperparameters for this model by varying the learning rate and the size of the fully connected layers and evaluating validation set performance, using one-hot encoded data as input.

### In silico directed evolution for COVID

Since our goal here was to find a sequence which binds tightly to all of the targets, we used the *in silico* directed evolution strategy with the conservative scheme described above under “Experiments on synthetic data from the Absolut! Database”. As discussed in Parkinson / Hard et al.^12^, even with *in silico* directed evolution, antibody sequence space is too large to search efficiently; thus, it is generally preferable to restrict the search space to some subset of the input sequence. This could be done in any of a number of ways as preferred by the end user – based on prior knowledge about which positions are most important, for example, or as in Parkinson / Hard et al.^12^ by selecting those positions which are most frequently mutated in tight binding sequences. We describe our process for selecting positions above under the Materials and Methods section under “Sequence processing”.

On each iteration of directed evolution, if the uncertainty for any prediction against any antigen is > the 80th percentile of variance values observed in the training set, the mutation was automatically rejected. This ensured that we did not consider scores less reliable than most scores assigned to the training set. We ran 5 *in silico* directed evolution experiments and harvested the accepted sequences.

Given our resources, our goal at this stage was to find the best <= 30 sequences to test experimentally. The best sequences were those with the highest scores (most likely to be tight binders), with the lowest variance (most confident predictions), the highest human-likeness (lowest risk of immunogenicity) and best predicted solubility. It is possible to incorporate all of these objectives into the *in silico* directed evolution procedure described above. Our preference, however, is to use the *in silico* process to generate new batches of likely tight binders as many times as needed, then filter these based on the other criteria to select a number of sequences compatible with resources available for experimental testing.

We therefore removed any sequences for which the smallest predicted score against any antigen was < 3.5 (the 95th percentile for scores in the training set), corresponding to a sequence which was roughly 30 times more abundant in the filtered bin than in the naive library. Our experimental capacity only allowed testing of up to 30 sequences. Therefore, we used a series cutoffs to select the most confident predictions. We first removed any sequences with human-likeness scores < −100 assigned by AntPack^29^ and further removed the 30% with the worst human-likeness scores. Next, we sorted sequences by the variance on the model-predicted score and discarded the worst 40%. This yielded 59 sequences; we then selected the 29 of these with the best score. The resulting sequences were experimentally tested as described above. Many of these filter cutoffs are arbitrary and can be adjusted if more (or fewer) sequences are desired for experimental testing.

### Generation and Screening of the 29 Member Delta-6 Library

The 29 dsDNA fragments used to create the 29-member library were purchased from IDT as eBlocks, followed by PCR amplification by Q5 hotstart DNA polymerase using “29 F/R” primers listed in supplemental table S7. The PCR products were concentrated with DNA Clean and Concentrator-5 kit (Zymo Research), purified by gel-extraction (1% agarose/Zymo Research Gel Recovery Kit), then double digested/ligated into the WT Delta-6 pCTCON2 vector by cutting out the WT section of the Delta-6 scFv with NheI-Hf & MluI-HF and ligating the mutant DNA with T4 DNA ligase (after treating the digested vector with rSAP (M0371S, NEB)). The plasmids were transformed into GC10 competent cells (Genesee Scientific 42-658) and sequenced by Sanger sequencing to verify the correct insert.

The 29 plasmids for the library were pooled together and transformed into EBY100 yeasts using the T2001 kit (Frozen-EZ Yeast Transformation II kit, Zymo Research). The WT Delta-6 scFv in pCTCON2 was separately transformed with the same kit. Transformants for each transformation (at least 7,000 colonies each) were scraped together into SDCAA citrate, grown to high density with shaking at 30C, then made into frozen stocks at −80C with freezing media (85% ddH20, 10% glycerol, 5% DMSO). To screen the 29-member library vs. WT Delta-6, the library and Delta-6 frozen yeasts stocks were thawed and added at 5 × 10^6^ viable cells each to 50 mL SDCAA-citrate and rotated at 30C/300RPM until the cultures got to high density, diluted 10x to OD600 = 0.5, rotated to high density again and induced once the OD600 reached 5. The cells were induced in SGCAA + ampicillin (100 μg/mL) at 20C, 300 RPM for 46h. The cells were then added at 100,000 cells per RBD tube (the concentration of the RBD in each tube depended on the K_D_ of that RBD to WT Delta-6). For the RBDs from WT SARS COV-2, Alpha, Delta, and Lambda variants, the concentration of the RBD was equal to the K_D_ of WT Delta-6 scFv. For Kappa, it was 160 nM (we estimated this was the K_D_ of Delta-6 to Kappa based on its binding curve), and 321 nM for the rest of the RBDs since the WT Delta-6 K_D_ was too weak to determine against them. Yeasts/RBDs were rotated in 1% BSA in PBS for 13.5 hr at RT, then 1:100 of 9B11 mouse anti-c-myc was added for 2h at RT. Afterwards, the yeasts were pelleted at 17,000xG/30s, washed with 100 uL cold PBS-BSA, then resuspended in 100 uL cold PBS-BSA with 1:100 each streptavidin-PE and goat anti-mouse AF647 for 20 minutes on ice in the dark with tapping. The tubes were pelleted high speed, washed with 100 uL cold PBS-BSA, then resuspened at 120 uL each for flow cytometry analysis. Solutions were analyzed at 50,000 events per tube on a BD FACSAria II cell sorter at the Moores Cancer Center UC San Diego. For each RBD, a gate was drawn just above the WT Delta-6 main population and the 29-member library was analyzed compared to the same gate to determine the % of library member cells occurring above the WT main group of cells.

### Determination of the Expression Level of Delta-63 and other scFv on the Yeast Surface

ySYNAGa yeasts were transformed with PmeI-digested scFv in the pSYNAGa plasmid using a Frozen-EZ Yeast Transformation II kit, (Zymo Research) and expression on the yeast surface was determined basically as described^64^, using a 1:50 dilution of fitc-mAb (CMYC-45F-Z (ICL)). The FITC mean fluorescent intensity was determined on a BD FACSAria II cell sorter at the Moores Cancer Center UC San Diego.

## SOFTWARE

Analysis and modeling was conducted using Python 3.10 with the PyTorch library version 2.3.1, the Numpy library version 1.24.4, the Scipy library version 1.13.1, the scikit-learn library version 1.5.0, and the AntPack library version 0.2.7. xGPR v0.2.0.5 was used for the COVID model, while the faster and more recent v0.4.6 was used for the synthetic data experiments. 123 was used as a random seed for model weight initialization, train-test splitting etc. xGPR in later versions (>0.4.0.01) is much faster and easier to install, so we recommend that xGPR > 0.4 should be used for future projects.

## STATISTICS AND REPRODUCIBILITY

No statistical method was used to predetermine sample size. When processing raw sequence data, unreliable sequence reads (using criteria detailed in the Supplementary Info section S1) were discarded before any further analysis or processing was conducted. These steps were taken to ensure that only reliable reads were used for analysis. No data was otherwise excluded from any subsequent analysis or model training. The test set for evaluating model performance was constructed by randomly selecting 20% of the assembled sequences and assigning these to test. The random partition was generated using the train_test_split function implemented in scikit-learn version 1.5.0 with a seed value of 123. When model performance was assessed using cross-validations, the cross-validation splits were generated by randomly partitioning the dataset into 5 splits of equal size using the KFold function in Python’s scikit-learn library version 1.5.0.

The final evaluation of model performance was conducted “blind” by generating predictions for sequences not present in our data and experimentally evaluating these predictions as described above.

## CODE AVAILABILITY

The code used to perform the analysis described in this paper is available at https://github.com/Wang-lab-UCSD/RESP2, together with the code for the resp_protein_toolkit which enables users to run a RESP optimization on their own data.

## Acknowledgements

This work is partially supported by NIH (R21AI158114 and R01AI150282). The naive scFv yeast display library was generously provided by the lab of Prof. Dane Wittrup (MIT), the pCTCON2 vector was from Addgene (plasmid #41843). The ySYNAGa yeast strain and pSYNAGa plasmid were provided by the lab of Prof. Eric Klavins (University of Washington).

## AUTHOR CONTRIBUTIONS STATEMENT

These authors contributed equally: Jonathan Parkinson, Ryan Hard. Jonathan Parkinson was responsible for all computational analysis and statistical modeling, Ryan Hard for all experiment design and execution. Young Su Ko was responsible for software testing and contributed ideas regarding the use of the SNGP uncertainty quantitation method. All authors contributed to planning and overall design of the study.

