## Supplementary Info for "RESP2: An uncertainty aware multi-target multi-property optimization AI pipeline for antibody discovery"

### Section S1. Sequence quality filtering procedure

Raw paired-end reads were processed by first eliminating any which contained N or for which the lowest phred quality score in a non-overlapping region fell below 8. Next, the overlapping regions of the paired-end reads were considered. In any case where the amino acid assigned based on one read did not match the other, the amino acid assigned based on the higher phred quality score was used, as long as the phred quality score was > 8. If none of these criteria were met the pair of reads was eliminated. Finally, any sequences containing stop codons or where the first 9 or last 9 amino acids had been modified were excluded, since in our approach we did not want to consider these regions for potential modification.

Next, any amino acid sequences present only once were excluded, since singletons are especially likely to be the result of sequencing errors and it is hard to make inferences about them (e.g. if a sequence is present only once, it is not clear if it is more or less abundant in the naive and filtered libraries). This filtering procedure yielded 9,058,099 amino acid sequences or 205,520 unique amino acid sequences with an average of 3.7 mutations per sequence as compared with the wild-type.

The frequency of each sequence in both the naive library and in the filtered bins was tabulated. Any sequences with a frequency of zero in the naive library but a nonzero frequency in filtered bins were set to have a frequency of 1 in the naive library by default, since all sequences present in the filtered bins must also originally have been present in the naive library; if they are absent it is merely due to insufficient sequencing depth. This correction was applied to 74,188 unique sequences.

### Section S2. Preprocessing of substitution matrices to yield substitution matrix distance.

Let $M_i$ be the *i*-th row of a preprocessed substitution matrix and let *R* be the raw substitution matrix. We would like the distance between any two amino acids to be $D_{ij} = \frac{max(R_{ii}, R_{jj}) - R_{ij}}{max(R_{ii}, R_{jj})}$ – i.e.

substitution matrix distance normalized by self-similarity. We would like to preprocess the raw substitution matrix *R* in such a way that $||M_i - M_j||^2 = D_{ij}$. This will then ensure that for the kernel $k(x, y) = e^{-\gamma||x-y||^2}$, the term in the exponent becomes the sum over $D_{ij}$ for all positions in the two sequences.

To achieve this, we first generate the matrix $D$ so that $D_{ij} = \frac{max(R_{ii}, R_{jj}) - R_{ij}}{max(R_{ii}, R_{jj})}$. This is a distance matrix. Diagonal values are set to be 0 since the distance of an amino acid from itself should be zero. We then convert *D* to a similarity matrix *S* by subtracting it from the *maximum* value in *D*. We then take the Cholesky decomposition *C* of *S* such that $S = CC^T$. It is easy to see that $C_i^T C_j = S_{ij}$ for rows *i* and *j* of *C*.

Now notice that $||M_i - M_j||^2 = M_i^T M_i + M_j^T M_j - 2M_i^T M_j$. If we encode each amino acid using the corresponding rows of *C*, we have that

$$||C_i - C_j||^2 = C_i^T C_i + C_j^T C_j - 2C_i^T C_j = S_{ii} + S_{jj} - 2S_{ij} = 2D_{max} - 2(D_{max} - D_{ij}) = 2D_{ij}$$

If we divide the rows of *C* by $\sqrt{2}$, the final result becomes $||C_i - C_j||^2 = D_{ij}$

So we can encode each amino acid using the corresponding row of *C* divided by $\sqrt{2}$. If we feed this into an appropriate kernel, we ensure that the term in the exponent of the kernel will be the substitution matrix distance, as desired.

**SUPPLEMENTARY TABLES**

**Table S1** Performance on synthetic data from the Absolut! Database for different model architectures, as measured using $R^2$, mean absolute error (MAE) and root mean squared error (RMSE) for predictions vs actual values. For these evaluations, we use a dataset of 500,000 sequences provided by Absolut which is randomly sampled from the bottom 99% of binders; this dataset is split into a train and test set with an 80-20% split and performance on the held-out test set is reported. All models use the CDRH3 data as input. All models use PFASUM encoding using a common substitution matrix derived from Pfam seed multiple sequence alignments (MSAs), which is similar to BLOSUM but tailored to the known sequence space for improved homology search results and MSA quality. Area under the calibration error curve (AUCE) is calculated for models which are uncertainty-aware; for all others it is omitted.

The following models are included:
*vBNN*: A three-layer variational Bayesian neural network used in RESP1.
*xGPR*: An approximate Gaussian process using the xGPR library with a FastConv1d kernel used in RESP2.

| Antigen Group | Antigen | Model | Test set $R^2$ | Test set MAE | Test set RMSE | Test set AUCE |
|---|---|---|---|---|---|---|
| Neuraminidase | 1NCA_N | xGPR | 0.891 | 1.379 | 1.836 | 0.096 |
| | 1NCA_N | vBNN | 0.316 | 3.8 | 4.592 | 0.395 |
| | 4QNP_A | xGPR | 0.918 | 1.111 | 1.6 | 0.024 |
| | 4QNP_A | vBNN | 0.618 | 2.861 | 3.464 | 0.417 |
| IL-2 | 4YUE_C | xGPR | 0.817 | 1.35 | 2.152 | 0.269 |
| | 4YUE_C | vBNN | 0.657 | 2.042 | 2.946 | 0.324 |
| | 5LQB_A | xGPR | 0.894 | 1.392 | 1.831 | 0.094 |
| | 5LQB_A | vBNN | 0.124 | 4.446 | 5.257 | 0.437 |
| Prion protein | 1TQB_A | xGPR | 0.966 | 0.759 | 1.085 | 0.162 |
| | 1TQB_A | vBNN | 0.752 | 2.372 | 2.937 | 0.357 |
| | 2W9E_A | xGPR | 0.818 | 1.322 | 2.245 | 0.025 |
| | 2W9E_A | vBNN | 0.447 | 2.908 | 3.913 | 0.344 |
| | 4H88_A | xGPR | 0.956 | 0.837 | 1.139 | 0.035 |
| | 4H88_A | vBNN | 0.761 | 2.157 | 2.668 | 0.318 |

| | | | | | | |
|---|---|---|---|---|---|---|
| Dust mite antigen | 3RVV_A | xGPR | 0.895 | 1.322 | 1.918 | 0.297 |
| | 3RVV_A | vBNN | 0.626 | 2.834 | 3.617 | 0.311 |
| | 4PP1_A | xGPR | 0.612 | 2.18 | 2.927 | 0.17 |
| | 4PP1_A | vBNN | 0.125 | 3.448 | 4.398 | 0.393 |
| Notch protein | 3L95_X | xGPR | 0.859 | 1.738 | 2.313 | 0.131 |
| | 3L95_X | vBNN | -0.115 | 5.602 | 6.499 | 0.456 |
| | 5CZV_A | xGPR | 0.903 | 1.365 | 1.831 | 0.289 |
| | 5CZV_A | vBNN | 0.622 | 2.951 | 3.607 | 0.363 |
| Tissue factor protein | 1JPS_T | xGPR | 0.967 | 0.862 | 1.132 | 0.241 |
| | 1JPS_T | vBNN | 0.691 | 2.832 | 3.455 | 0.393 |
| | 4M7L_T | xGPR | 0.888 | 1.129 | 1.775 | 0.032 |
| | 4M7L_T | vBNN | 0.665 | 2.387 | 3.07 | 0.361 |

**Table S2** Success rates with bootstrapped confidence intervals (nonparametric bootstrap) for different candidate generation techniques on the Absolut! database. Success here is defined as an instance where a sequence generated as a candidate by an algorithm achieves a binding score tighter than any training set sequence for all antigens in the group.

| Target group | Candidate generation approach | Number of unique candidates generated | Success rate (95% CI) |
|---|---|---|---|
| Dust mite antigen | Evodiff-80 | 1000 | 0.0 (0 - 0) |
| IL-2 | Evodiff-80 | 1000 | 0.9 (0.4 - 1.5) |
| Neuraminidase | Evodiff-80 | 999 | 1.0 (0.4 - 1.7) |
| Notch protein | Evodiff-80 | 1000 | 0.5 (0.1 - 0.9) |
| Prion protein | Evodiff-80 | 1000 | 0.1 (0 - 0.3) |

| | | | |
|---|---|---|---|
| Tissue factor protein | Evodiff-80 | 999 | 0.6 (0.2 - 1.2) |
| Dust mite antigen | Evodiff-90 | 1000 | 0.5 (0.1 - 1.0) |
| IL-2 | Evodiff-90 | 1000 | 1.3 (0.7 - 2.0) |
| Neuraminidase | Evodiff-90 | 1000 | 1.2 (0.6 - 2.0) |
| Notch protein | Evodiff-90 | 1000 | 0.9 (0.4 - 1.5) |
| Prion protein | Evodiff-90 | 1000 | 0.0 (0 - 0) |
| Tissue factor protein | Evodiff-90 | 1000 | 1.8 |
| Dust mite antigen | Protein-MPNN | 629 | 0.0 (0 - 0) |
| IL-2 | Protein-MPNN | 373 | 0.0 (0 - 0) |
| Neuraminidase | Protein-MPNN | 153 | 0.0 (0 - 0) |
| Notch protein | Protein-MPNN | 76 | 0.0 (0 - 0) |
| Prion protein | Protein-MPNN | 53 | 0.0 (0 - 0) |
| Tissue factor protein | Protein-MPNN | 184 | 0.0 (0 - 0) |
| Dust mite antigen | RESP | 234 | 88.0 (83.8 - 91.9) |
| IL-2 | RESP | 547 | 89.8 (87.4 - 92.1) |
| Neuraminidase | RESP | 1031 | 99.5 (99.0 - 99.9) |
| Notch protein | RESP | 729 | 91.1 (89.0 - 93.0) |
| Prion protein | RESP | 79 | 100.0 (100 - 100) |
| Tissue factor protein | RESP | 289 | 84.8 (80.3 - 88.6) |

**Table S3** Performance on the test set for Group A antigens with different encoding schemes and model architectures, as measured using $R^2$, mean absolute error (MAE) and root mean squared error (RMSE) for predictions vs actual values. The 95% CI is calculated by bootstrapping and is in parentheses.

| Encoding type for antibody | Encoding type for antigen | Model architecture | Uncertainty-aware? | Test set R^2 (95% CI) | Test set MAE (95% CI) | Test set RMSE (95% CI) |
|---|---|---|---|---|---|---|
| One-hot | One-hot | Linear | Yes | 0.596 (0.583 - 0.609) | 0.571 (0.56 - 0.581) | 0.965 (0.944 - 0.986) |

| | | | | | | |
|---|---|---|---|---|---|---|
| One-hot | One-hot | Variational Bayesian NN (RESP architecture) | Yes | 0.722 (0.709 - 0.734) | 0.38 (0.371 - 0.390) | 0.801 (0.778 - 0.824) |
| One-hot | One-hot | xGPR, RBF kernel | Yes | 0.765 (0.756 - 0.774) | 0.425 (0.417 - 0.433) | 0.737 (0.719 - 0.755) |
| One-hot | One-hot | ByteNet | No | 0.767 (0.756 - 0.778) | 0.307 (0.297 - 0.315) | 0.734 (0.712 - 0.755) |
| One-hot | One-hot | ByteNet, last layer GP | Yes | 0.747 (0.737 - 0.757) | 0.409 (0.4 - 0.417) | 0.764 (0.745 - 0.784) |
| One-hot | One-hot | xGPR, FastConv1d kernel | Yes | 0.759 (0.749 - 0.769) | 0.426 (0.418 - 0.435) | 0.746 (0.727 - 0.766) |
| AbLang embedding | ESM embedding (not averaged) | Linear | Yes | 0.767 (0.757 - 0.776) | 0.409 (0.400 - 0.417) | 0.734 (0.716 - 0.753) |
| AbLang embedding | ESM embedding (not averaged) | xGPR, RBF kernel, different lengthscale for antibody and antigen | Yes | 0.744 (0.734 - 0.754) | 0.433 (0.424 - 0.441) | 0.769 (0.750 - 0.790) |
| AbLang embedding | ESM embedding (not averaged) | Variational Bayesian NN (RESP architecture) | Yes | 0.668 (0.656 - 0.681) | 0.401 (0.389 - 0.412) | 0.875 (0.851 - 0.899) |
| AbLang embedding | ESM embedding (not averaged) | ByteNet | No | 0.781 (0.770 - 0.792) | 0.308 (0.299 - 0.316) | 0.721 (0.700 - 0.742) |
| AbLang embedding | ESM embedding (not averaged) | ByteNet, last layer GP | Yes | 0.762 (0.751 - 0.773) | 0.339 (0.330 - 0.348) | 0.733 (0.712 - 0.754) |
| AbLang embedding | ESM embedding (averaged) | xGPR, FastConv1d kernel | Yes | 0.753 (0.743 - 0.763) | 0.436 (0.427 - 0.444) | 0.755 (0.736 - 0.774) |
| One-hot | ESM embedding, averaged over antigen sequence | xGPR, RBF kernel | Yes | 0.763 (0.754 - 0.772) | 0.426 (0.417 - 0.434) | 0.740 (0.721 - 0.759) |
| One-hot | ESM embedding, averaged over antigen sequence | Variational Bayesian NN (RESP architecture) | Yes | 0.483 (0.470 - 0.495) | 0.569 (0.557 - 0.582) | 1.093 (1.068 - 1.116) |
| One-hot | ESM embedding, averaged over antigen sequence | Linear | Yes | 0.596 (0.583 - 0.609) | 0.57 (0.559 - 0.581) | 0.965 (0.944 - 0.986) |

| | | | | | | |
|---|---|---|---|---|---|---|
| One-hot | ESM embedding, averaged over antigen sequence | xGPR, FastConv1d kernel | Yes | 0.759 (0.749 - 0.769) | 0.426 (0.418 - 0.434) | 0.746 (0.727 - 0.766) |
| PFASUM matrix | PFASUM matrix | Linear | Yes | 0.596 (0.583 - 0.609) | 0.570 (0.560 - 0.581) | 0.965 (0.944 - 0.986) |
| PFASUM matrix | PFASUM matrix | Variational Bayesian NN (RESP architecture) | Yes | 0.719 (0.708 - 0.730) | 0.45 (0.441 - 0.459) | 0.806 (0.785 - 0.825) |
| PFASUM matrix | PFASUM matrix | xGPR, RBF kernel | Yes | 0.766 (0.756 - 0.774) | 0.424 (0.416 - 0.433) | 0.736 (0.717 - 0.754) |
| PFASUM matrix | PFASUM matrix | ByteNet | No | 0.775 (0.764 - 0.786) | 0.314 (0.305 - 0.323) | 0.720 (0.700 - 0.741) |
| PFASUM matrix | PFASUM matrix | ByteNet, last layer GP | Yes | 0.762 (0.751 - 0.773) | 0.367 (0.358 - 0.375) | 0.741 (0.721 - 0.762) |
| PFASUM matrix | PFASUM matrix | xGPR, FastConv1d kernel | Yes | 0.764 (0.754 - 0.773) | 0.420 (0.412 - 0.429) | 0.739 (0.720 - 0.759) |
| RESP 1.0 autoencoder | One-hot encoding | Linear | Yes | 0.577 (0.563 - 0.589) | 0.602 (0.591 - 0.613) | 0.989 (0.968 - 1.009) |
| RESP 1.0 autoencoder | One-hot encoding | Variational Bayesian NN (RESP architecture) | Yes | 0.697 (0.686 - 0.709) | 0.431 (0.421 - 0.442) | 0.836 (0.814 - 0.858) |
| RESP 1.0 autoencoder | One-hot encoding | xGPR, RBF kernel | Yes | 0.743 (0.733 - 0.753) | 0.445 (0.436 - 0.454) | 0.771 (0.752 - 0.789) |
| RESP 1.0 autoencoder | One-hot encoding | ByteNet | No | 0.775 (0.764 - 0.787) | 0.323 (0.314 - 0.332) | 0.720 (0.699 - 0.741) |
| RESP 1.0 autoencoder | One-hot encoding | ByteNet, last layer GP | Yes | 0.765 (0.754 - 0.775) | 0.383 (0.374 - 0.391) | 0.737 (0.717 - 0.759) |
| RESP 1.0 autoencoder | One-hot encoding | xGPR, FastConv1d kernel | Yes | 0.730 (0.719 - 0.741) | 0.445 (0.436 - 0.454) | 0.790 (0.770 - 0.809) |

**Table S4** Area under the calibration error curve (AUCE) for different model architectures.

| **Encoding type for antibody** | **Encoding type for antigen** | **Model architecture** | **AUCE on test set** |
|---|---|---|---|
| One-hot | One-hot | Variational Bayesian NN (RESP 1.0 architecture) | 0.282 |
| One-hot | One-hot | ByteNet, last layer GP | 0.308 |

| | | | |
|---|---|---|---|
| One-hot | One-hot | xGPR, FastConv1d kernel | 0.306 |
| One-hot | One-hot | xGPR, RBF kernel | 0.241 |
| One-hot | One-hot | Linear | 0.396 |
| PFASUM matrix | PFASUM matrix | Linear | 0.376 |
| PFASUM matrix | PFASUM matrix | Variational Bayesian NN (RESP architecture) | 0.316 |
| PFASUM matrix | PFASUM matrix | xGPR, RBF kernel | 0.293 |
| PFASUM matrix | PFASUM matrix | ByteNet, last layer GP | 0.330 |
| PFASUM matrix | PFASUM matrix | xGPR, FastConv1d kernel | 0.291 |
| RESP 1.0 autoencoder | One-hot encoding | Linear | 0.162 |
| RESP 1.0 autoencoder | One-hot encoding | Variational Bayesian NN (RESP architecture) | 0.277 |
| RESP 1.0 autoencoder | One-hot encoding | xGPR, RBF kernel | 0.234 |
| RESP 1.0 autoencoder | One-hot encoding | ByteNet, last layer GP | 0.320 |
| RESP 1.0 autoencoder | One-hot encoding | xGPR, FastConv1d kernel | 0.075 |
| One-hot | ESM embedding, averaged over antigen sequence | xGPR, RBF kernel | 0.239 |
| One-hot | ESM embedding, averaged over antigen sequence | Variational Bayesian NN (RESP architecture) | 0.295 |
| One-hot | ESM embedding, averaged over antigen sequence | Linear | 0.396 |
| AbLang embedding | ESM embedding (not averaged) | Linear | 0.495 |
| AbLang embedding | ESM embedding (not averaged) | xGPR, RBF kernel, different lengthscale for antibody and antigen | 0.326 |
| AbLang embedding | ESM embedding (not averaged) | Variational Bayesian NN (RESP architecture) | 0.305 |

| | | | |
|---|---|---|---|
| AbLang embedding | ESM embedding (not averaged) | ByteNet, last layer GP | 0.316 |
| AbLang embedding | ESM embedding (averaged) | xGPR, FastConv1d kernel | 0.191 |

**Table S5:** The target protein groups from the Absolut! database used for the Absolut! synthetic data experiments.

| Target protein group | PDB code for antigen complex structure | Maximum pairwise percent identity to any other antigen in same group |
|---|---|---|
| Neuraminidase | 1NCA_N | 55.96 |
| | 4QNP_A | 55.96 |
| IL-2 | 4YUE_C | 63.8 |
| | 5LQB_A | 63.8 |
| Prion protein | 1TQB_A | 91.2 |
| | 2W9E_A | 90.2 |
| | 4H88_A | 91.2 |
| Dust mite antigen | 3RVV_A | 82.4 |
| | 4PP1_A | 82.4 |
| Notch protein | 3L95_X | 57.8 |
| | 3CZV_A | 57.8 |
| Tissue factor protein | 1JPS_T | 97.2 |
| | 4M7L_T | 97.2 |

**Table S6**: The variable heavy chain amino acid sequences of the WT Delta-6 scFv and 29-member scFv library (Delta-63 is mutant 26).

| scFv | HC Mutations | scFv HC sequence |
|---|---|---|
| Delta-6 | None (WT) | EVQLLESGGGLVQPGGTLRLSCAASGFIVSSNYMSWVRQAPGKGLEWVSLVYPGGSTYYADSVKGRFTVSRDNSKNTLYLQMNSLRAEDMAVYYCARDLPSGVDAVDAFDIWGQGTMVTVSS |
| Mutant 0 | G26V_F27I | EVQLLESGGGLVQPGGTLRLSCAASVIIVSSNYMSWVRQAPGKGLEWVSLVYPGGSTYYADSVKGRFTVSRDNSKNTLYLQMNSLRAEDMAVYYCARDLPSGVDAVDAFDIWGQGTMVTVSS |
| Mutant 1 | F27I_V103I | EVQLLESGGGLVQPGGTLRLSCAASGIIVSSNYMSWVRQAPGKGLEWVSLVYPGGSTYYADSVKGRFTVSRDNSKNTLYLQMNSLRAEDMAVYYCARDLPSGIDAVDAFDIWGQGTMVTVSS |
| Mutant 2 | F27L_N73M | EVQLLESGGGLVQPGGTLRLSCAASGLIVSSNYMSWVRQAPGKGLEWVSLVYPGGSTYYADSVKGRFTVSRDMSKNTLYLQMNSLRAEDMAVYYCARDLPSGVDAVDAFDIWGQGTMVTVSS |
| Mutant 3 | G26E_F27V_N73S_V103I | EVQLLESGGGLVQPGGTLRLSCAASEVIVSSNYMSWVRQAPGKGLEWVSLVYPGGSTYYADSVKGRFTVSRDSSKNTLYLQMNSLRAEDMAVYYCARDLPSGIDAVDAFDIWGQGTMVTVSS |
| Mutant 4 | G26E_F27I_N73S_V103I | EVQLLESGGGLVQPGGTLRLSCAASEIIVSSNYMSWVRQAPGKGLEWVSLVYPGGSTYYADSVKGRFTVSRDSSKNTLYLQMNSLRAEDMAVYYCARDLPSGIDAVDAFDIWGQGTMVTVSS |
| Mutant 5 | G26E_F27V_N73D_V103I | EVQLLESGGGLVQPGGTLRLSCAASEVIVSSNYMSWVRQAPGKGLEWVSLVYPGGSTYYADSVKGRFTVSRDDSKNTLYLQMNSLRAEDMAVYYCARDLPSGIDAVDAFDIWGQGTMVTVSS |
| Mutant 6 | G26E_F27M_V103I | EVQLLESGGGLVQPGGTLRLSCAASEMIVSSNYMSWVRQAPGKGLEWVSLVYPGGSTYYADSVKGRFTVSRDNSKNTLYLQMNSLRAEDMAVYYCARDLPSGIDAVDAFDIWGQGTMVTVSS |
| Mutant 7 | G26E_F27I_V103I | EVQLLESGGGLVQPGGTLRLSCAASEIIVSSNYMSWVRQAPGKGLEWVSLVYPGGSTYYADSVKGRFTVSRDNSKNTLYLQMNSLRAEDMAVYYCARDLPSGIDAVDAFDIWGQGTMVTVSS |
| Mutant 8 | G26E_F27V | EVQLLESGGGLVQPGGTLRLSCAASEVIVSSNYMSWVRQAPGKGLEWVSLVYPGGSTYYADSVKGRFTVSRDNSKNTLYLQMNSLRAEDMAVYYCARDLPSGVDAVDAFDIWGQGTMVTVSS |
| Mutant 9 | G26E_F27L_V103N | EVQLLESGGGLVQPGGTLRLSCAASELIVSSNYMSWVRQAPGKGLEWVSLVYPGGSTYYADSVKGRFTVSRDNSKNTLYLQMNSLRAEDMAVYYCARDLPSGNDAVDAFDIWGQGTMVTVSS |

| | | |
|---|---|---|
| Mutant 10 | F27L_N73D_V103I | EVQLLESGGGLVQPGGTLRLSCAASGLIVSSNYMSWVRQAPGKGLEWVSLVYPGGSTYYADSVKGRFTVSRDDSKNTLYLQMNSLRAEDMAVYYCARDLPSGIDAVDAFDIWGQGTMVTVSS |
| Mutant 11 | F27L_V29A_N73D_V103I | EVQLLESGGGLVQPGGTLRLSCAASGLIASSNYMSWVRQAPGKGLEWVSLVYPGGSTYYADSVKGRFTVSRDDSKNTLYLQMNSLRAEDMAVYYCARDLPSGIDAVDAFDIWGQGTMVTVSS |
| Mutant 12 | F27L_V29A_V103I | EVQLLESGGGLVQPGGTLRLSCAASGLIASSNYMSWVRQAPGKGLEWVSLVYPGGSTYYADSVKGRFTVSRDNSKNTLYLQMNSLRAEDMAVYYCARDLPSGIDAVDAFDIWGQGTMVTVSS |
| Mutant 13 | F27L_V29D_V103I | EVQLLESGGGLVQPGGTLRLSCAASGLIDSSNYMSWVRQAPGKGLEWVSLVYPGGSTYYADSVKGRFTVSRDNSKNTLYLQMNSLRAEDMAVYYCARDLPSGIDAVDAFDIWGQGTMVTVSS |
| Mutant 14 | F27L_V29E_V103I | EVQLLESGGGLVQPGGTLRLSCAASGLIESSNYMSWVRQAPGKGLEWVSLVYPGGSTYYADSVKGRFTVSRDNSKNTLYLQMNSLRAEDMAVYYCARDLPSGIDAVDAFDIWGQGTMVTVSS |
| Mutant 15 | G26E_F27L_N73D_V103I | EVQLLESGGGLVQPGGTLRLSCAASELIVSSNYMSWVRQAPGKGLEWVSLVYPGGSTYYADSVKGRFTVSRDDSKNTLYLQMNSLRAEDMAVYYCARDLPSGIDAVDAFDIWGQGTMVTVSS |
| Mutant 16 | G26E_F27L_N73D | EVQLLESGGGLVQPGGTLRLSCAASELIVSSNYMSWVRQAPGKGLEWVSLVYPGGSTYYADSVKGRFTVSRDDSKNTLYLQMNSLRAEDMAVYYCARDLPSGVDAVDAFDIWGQGTMVTVSS |
| Mutant 17 | G26E_F27I_N73I | EVQLLESGGGLVQPGGTLRLSCAASEIIVSSNYMSWVRQAPGKGLEWVSLVYPGGSTYYADSVKGRFTVSRDISKNTLYLQMNSLRAEDMAVYYCARDLPSGVDAVDAFDIWGQGTMVTVSS |
| Mutant 18 | G26E_F27I_N73S | EVQLLESGGGLVQPGGTLRLSCAASEIIVSSNYMSWVRQAPGKGLEWVSLVYPGGSTYYADSVKGRFTVSRDSSKNTLYLQMNSLRAEDMAVYYCARDLPSGVDAVDAFDIWGQGTMVTVSS |
| Mutant 19 | G26E_F27L_N73S_V103S | EVQLLESGGGLVQPGGTLRLSCAASELIVSSNYMSWVRQAPGKGLEWVSLVYPGGSTYYADSVKGRFTVSRDSSKNTLYLQMNSLRAEDMAVYYCARDLPSGSDAVDAFDIWGQGTMVTVSS |
| Mutant 20 | G26E_F27L_N73S | EVQLLESGGGLVQPGGTLRLSCAASELIVSSNYMSWVRQAPGKGLEWVSLVYPGGSTYYADSVKGRFTVSRDSSKNTLYLQMNSLRAEDMAVYYCARDLPSGVDAVDAFDIWGQGTMVTVSS |
| Mutant 21 | G26E_F27I | EVQLLESGGGLVQPGGTLRLSCAASEIIVSSNYMSWVRQAPGKGLEWVSLVYPGGSTYYADSVKGRFTVSRDNSKNTLYLQMNSLRAEDMAVYYCARDLPSGVDAVDAFDIWGQGTMVTVSS |
| Mutant 22 | G26E_F27L | EVQLLESGGGLVQPGGTLRLSCAASELIVSSNYMSWVRQAPGKGLEWVSLVYPGGSTYYADSVKGRFTVSRDNSKNTLYLQMNSLRAEDMAVYYCARDLPSGVDAVDAFDIWGQGTMVTVSS |

| | | |
|---|---|---|
| Mutant 23 | G26E_F27I_N73T | EVQLLESGGGLVQPGGTLRLSCAASEIIVSSNYMSWVRQAPGKGLEWVSLVYPGGSTYYADSVKGRFTVSRDTSKNTLYLQMNSLRAEDMAVYYCARDLPSGVDAVDAFDIWGQGTMVTVSS |
| Mutant 24 | F27I_N73D_V103I | EVQLLESGGGLVQPGGTLRLSCAASGIIVSSNYMSWVRQAPGKGLEWVSLVYPGGSTYYADSVKGRFTVSRDDSKNTLYLQMNSLRAEDMAVYYCARDLPSGIDAVDAFDIWGQGTMVTVSS |
| Mutant 25 | G26E_F27L_V103I | EVQLLESGGGLVQPGGTLRLSCAASELIVSSNYMSWVRQAPGKGLEWVSLVYPGGSTYYADSVKGRFTVSRDNSKNTLYLQMNSLRAEDMAVYYCARDLPSGIDAVDAFDIWGQGTMVTVSS |
| Mutant 26 (Delta-63) | G26E_F27L_N73E_V103I | EVQLLESGGGLVQPGGTLRLSCAASELIVSSNYMSWVRQAPGKGLEWVSLVYPGGSTYYADSVKGRFTVSRDESKNTLYLQMNSLRAEDMAVYYCARDLPSGIDAVDAFDIWGQGTMVTVSS |
| Mutant 27 | F27L_N73I_V103I | EVQLLESGGGLVQPGGTLRLSCAASGLIVSSNYMSWVRQAPGKGLEWVSLVYPGGSTYYADSVKGRFTVSRDISKNTLYLQMNSLRAEDMAVYYCARDLPSGIDAVDAFDIWGQGTMVTVSS |
| Mutant 28 | F27I_N73I_V103I | EVQLLESGGGLVQPGGTLRLSCAASGIIVSSNYMSWVRQAPGKGLEWVSLVYPGGSTYYADSVKGRFTVSRDISKNTLYLQMNSLRAEDMAVYYCARDLPSGIDAVDAFDIWGQGTMVTVSS |

**Table S7**. Primers used to generate the MiSeq, random Delta-6, and 29-member libraries. Primers were purchased from IDT as standard desalted oligos.

| Primer Name | Primer Sequence (5'-3') |
|---|---|
| Delta-6 NGS F | TCGTCGGCAGCGTCAGATGTGTATAAGAGACAGTCGGCTAGCGAG |
| Delta-6 NGS R | GTCTCGTGGGCTCGGAGATGTGTATAAGAGACAGAGAATCCCTGAACTGAC |
| Ad1.1 | AATGATACGGCGACCACCGAGATCTACACTAGATCGCTCGTCGGCAGCGTCAGATGTG |
| Ad2.1 | CAAGCAGAAGACGGCATACGAGATTCGCCTTAGTCTCGTGGGCTCGGAGATGT |
| Ad2.2 | CAAGCAGAAGACGGCATACGAGATCTAGTACGGTCTCGTGGGCTCGGAGATGT |
| Ad2.3 | CAAGCAGAAGACGGCATACGAGATTTCTGCCTGTCTCGTGGGCTCGGAGATGT |
| Ad2.4 | CAAGCAGAAGACGGCATACGAGATGCTCAGGAGTCTCGTGGGCTCGGAGATGT |
| Ad2.5 | CAAGCAGAAGACGGCATACGAGATAGGAGTCCGTCTCGTGGGCTCGGAGATGT |
| Ad2.6 | CAAGCAGAAGACGGCATACGAGATCATGCCTAGTCTCGTGGGCTCGGAGATGT |
| Ad2.7 | CAAGCAGAAGACGGCATACGAGATGTAGAGAGGTCTCGTGGGCTCGGAGATGT |

| | |
|---|---|
| Ad2.8 | CAAGCAGAAGACGGCATACGAGATCCTCTCTGGTCTCGTGGGCTCGGAGATGT |
| Ad2.9 | CAAGCAGAAGACGGCATACGAGATAGCGTAGCGTCTCGTGGGCTCGGAGATGT |
| Ad2.10 | CAAGCAGAAGACGGCATACGAGATCAGCCTCGGTCTCGTGGGCTCGGAGATGT |
| Ad2.11 | CAAGCAGAAGACGGCATACGAGATTGCCTCTTGTCTCGTGGGCTCGGAGATGT |
| Ad2.12 | CAAGCAGAAGACGGCATACGAGATTCCTCTACGTCTCGTGGGCTCGGAGATGT |
| Ad2.13 | CAAGCAGAAGACGGCATACGAGATATCACGACGTCTCGTGGGCTCGGAGATGT |
| Ad2.14 | CAAGCAGAAGACGGCATACGAGATACAGTGGTGTCTCGTGGGCTCGGAGATGT |
| Ad2.15 | CAAGCAGAAGACGGCATACGAGATCAGATCCAGTCTCGTGGGCTCGGAGATGT |
| Ad2.16 | CAAGCAGAAGACGGCATACGAGATACAAACGGGTCTCGTGGGCTCGGAGATGT |
| Ad2.17 | CAAGCAGAAGACGGCATACGAGATACCCAGCAGTCTCGTGGGCTCGGAGATGT |
| Ad2.18 | CAAGCAGAAGACGGCATACGAGATAACCCCTCGTCTCGTGGGCTCGGAGATGT |
| Ad2.19 | CAAGCAGAAGACGGCATACGAGATCCCAACCTGTCTCGTGGGCTCGGAGATGT |
| Ad2.20 | CAAGCAGAAGACGGCATACGAGATCACCACACGTCTCGTGGGCTCGGAGATGT |
| Ad2.21 | CAAGCAGAAGACGGCATACGAGATGAAACCCAGTCTCGTGGGCTCGGAGATGT |
| Ad2.22 | CAAGCAGAAGACGGCATACGAGATTGTGACCAGTCTCGTGGGCTCGGAGATGT |
| Ad2.23 | CAAGCAGAAGACGGCATACGAGATAGGGTCAAGTCTCGTGGGCTCGGAGATGT |
| 29 F | TCAGCTAGCGAGGTCCAACTTC |
| 29 R | TATCGAACGCGTCAACGGCATC |
| D6 ePCR F | ACTTCTAGAATCTGGTGG |
| D6 ePCR R | CGTCACCATAGTACCTTG |
| D6F | CGGCTAGCGAGGTCCAACTTCTAGAATCTGGTGG |
| D6R | CACTTCCTAGAATCCCTGAACTGACCGTCACCATAGTACCTTG |
| D6 Linear F | GTTCAGGGATTCTAGGAAG |
| D6 Linear R | GAAGTTGGACCTCGCTAG |

**Table S8**. DNA sequences of mutants amplified by PCR for insertion into WT Delta-6 pCTCON2 vector.

| scFv | HC Mutations | eBlock Sequence |
|---|---|---|
| Mutant 0 | G26V_F27I | TCGGCTAGCGAGGTCCAACTTCTAGAATCTGGTGGGGGGCTTGTCCAGCCTGGTGGCACTTTGAGGCTGTCTTGTGCTGCCTCAGTTAT<br>TATCGTAAGTAGTAATTACATGAGTTGGGTGCGTCAGGCACCCGGAAAAGGGCTTGAATGGGTTAGTTTAGTTTACCCAGGTGGTAGCAC<br>TTACTATGCCGACTCTGTCAAAGGTAGGTTCACCGTTAGTCGTGATAATTCTAAAAACACTTTATACCTGCAAATGAATTCACTACGTGCTG<br>AAGACATGGCTGTTTATTATTGCGCACGTGACCTTCCATCCGGCGTTGATGCCGTTGACGCGTTCGATA |
| Mutant 1 | F27I_V103I | TCGGCTAGCGAGGTCCAACTTCTAGAATCTGGTGGGGGGCTTGTCCAGCCTGGTGGCACTTTGAGGCTGTCTTGTGCTGCCTCAGGGA<br>TTATCGTAAGTAGTAATTACATGAGTTGGGTGCGTCAGGCACCCGGAAAAGGGCTTGAATGGGTTAGTTTAGTTTACCCAGGTGGTAGCA<br>CTTACTATGCCGACTCTGTCAAAGGTAGGTTCACCGTTAGTCGTGATAATTCTAAAAACACTTTATACCTGCAAATGAATTCACTACGTGCT<br>GAAGACATGGCTGTTTATTATTGCGCACGTGACCTTCCATCCGGCATTGATGCCGTTGACGCGTTCGATA |
| Mutant 2 | F27L_N73M | TCGGCTAGCGAGGTCCAACTTCTAGAATCTGGTGGGGGGCTTGTCCAGCCTGGTGGCACTTTGAGGCTGTCTTGTGCTGCCTCAGGGT<br>TGATCGTAAGTAGTAATTACATGAGTTGGGTGCGTCAGGCACCCGGAAAAGGGCTTGAATGGGTTAGTTTAGTTTACCCAGGTGGTAGCA<br>CTTACTATGCCGACTCTGTCAAAGGTAGGTTCACCGTTAGTCGTGATATGTCTAAAAACACTTTATACCTGCAAATGAATTCACTACGTGCT<br>GAAGACATGGCTGTTTATTATTGCGCACGTGACCTTCCATCCGGCGTTGATGCCGTTGACGCGTTCGATA |
| Mutant 3 | G26E_F27V_N73S_V103I | TCGGCTAGCGAGGTCCAACTTCTAGAATCTGGTGGGGGGCTTGTCCAGCCTGGTGGCACTTTGAGGCTGTCTTGTGCTGCCTCAGAAG<br>TTATCGTAAGTAGTAATTACATGAGTTGGGTGCGTCAGGCACCCGGAAAAGGGCTTGAATGGGTTAGTTTAGTTTACCCAGGTGGTAGCA<br>CTTACTATGCCGACTCTGTCAAAGGTAGGTTCACCGTTAGTCGTGATTCTTCTAAAAACACTTTATACCTGCAAATGAATTCACTACGTGCT<br>GAAGACATGGCTGTTTATTATTGCGCACGTGACCTTCCATCCGGCATTGATGCCGTTGACGCGTTCGATA |
| Mutant 4 | G26E_F27I_N73S_V103I | TCGGCTAGCGAGGTCCAACTTCTAGAATCTGGTGGGGGGCTTGTCCAGCCTGGTGGCACTTTGAGGCTGTCTTGTGCTGCCTCAGAAAT<br>TATCGTAAGTAGTAATTACATGAGTTGGGTGCGTCAGGCACCCGGAAAAGGGCTTGAATGGGTTAGTTTAGTTTACCCAGGTGGTAGCAC<br>TTACTATGCCGACTCTGTCAAAGGTAGGTTCACCGTTAGTCGTGATTCTTCTAAAAACACTTTATACCTGCAAATGAATTCACTACGTGCTG<br>AAGACATGGCTGTTTATTATTGCGCACGTGACCTTCCATCCGGCATTGATGCCGTTGACGCGTTCGATA |
| Mutant 5 | G26E_F27V_N73D_V103I | TCGGCTAGCGAGGTCCAACTTCTAGAATCTGGTGGGGGGCTTGTCCAGCCTGGTGGCACTTTGAGGCTGTCTTGTGCTGCCTCAGAAG<br>TTATCGTAAGTAGTAATTACATGAGTTGGGTGCGTCAGGCACCCGGAAAAGGGCTTGAATGGGTTAGTTTAGTTTACCCAGGTGGTAGCA<br>CTTACTATGCCGACTCTGTCAAAGGTAGGTTCACCGTTAGTCGTGATGATTCTAAAAACACTTTATACCTGCAAATGAATTCACTACGTGCT<br>GAAGACATGGCTGTTTATTATTGCGCACGTGACCTTCCATCCGGCATTGATGCCGTTGACGCGTTCGATA |
| Mutant 6 | G26E_F27M_V103I | TCGGCTAGCGAGGTCCAACTTCTAGAATCTGGTGGGGGGCTTGTCCAGCCTGGTGGCACTTTGAGGCTGTCTTGTGCTGCCTCAGAAAT<br>GATCGTAAGTAGTAATTACATGAGTTGGGTGCGTCAGGCACCCGGAAAAGGGCTTGAATGGGTTAGTTTAGTTTACCCAGGTGGTAGCAC<br>TTACTATGCCGACTCTGTCAAAGGTAGGTTCACCGTTAGTCGTGATAATTCTAAAAACACTTTATACCTGCAAATGAATTCACTACGTGCTG<br>AAGACATGGCTGTTTATTATTGCGCACGTGACCTTCCATCCGGCATTGATGCCGTTGACGCGTTCGATA |
| Mutant 7 | G26E_F27I_V103I | TCGGCTAGCGAGGTCCAACTTCTAGAATCTGGTGGGGGGCTTGTCCAGCCTGGTGGCACTTTGAGGCTGTCTTGTGCTGCCTCAGAAAT<br>TATCGTAAGTAGTAATTACATGAGTTGGGTGCGTCAGGCACCCGGAAAAGGGCTTGAATGGGTTAGTTTAGTTTACCCAGGTGGTAGCAC<br>TTACTATGCCGACTCTGTCAAAGGTAGGTTCACCGTTAGTCGTGATAATTCTAAAAACACTTTATACCTGCAAATGAATTCACTACGTGCTG<br>AAGACATGGCTGTTTATTATTGCGCACGTGACCTTCCATCCGGCATTGATGCCGTTGACGCGTTCGATA |
| Mutant 8 | G26E_F27V | TCGGCTAGCGAGGTCCAACTTCTAGAATCTGGTGGGGGGCTTGTCCAGCCTGGTGGCACTTTGAGGCTGTCTTGTGCTGCCTCAGAAG<br>TTATCGTAAGTAGTAATTACATGAGTTGGGTGCGTCAGGCACCCGGAAAAGGGCTTGAATGGGTTAGTTTAGTTTACCCAGGTGGTAGCA<br>CTTACTATGCCGACTCTGTCAAAGGTAGGTTCACCGTTAGTCGTGATAATTCTAAAAACACTTTATACCTGCAAATGAATTCACTACGTGCT<br>GAAGACATGGCTGTTTATTATTGCGCACGTGACCTTCCATCCGGCGTTGATGCCGTTGACGCGTTCGATA |
| Mutant 9 | G26E_F27L_V103N | TCGGCTAGCGAGGTCCAACTTCTAGAATCTGGTGGGGGGCTTGTCCAGCCTGGTGGCACTTTGAGGCTGTCTTGTGCTGCCTCAGAATT<br>GATCGTAAGTAGTAATTACATGAGTTGGGTGCGTCAGGCACCCGGAAAAGGGCTTGAATGGGTTAGTTTAGTTTACCCAGGTGGTAGCAC<br>TTACTATGCCGACTCTGTCAAAGGTAGGTTCACCGTTAGTCGTGATAATTCTAAAAACACTTTATACCTGCAAATGAATTCACTACGTGCTG<br>AAGACATGGCTGTTTATTATTGCGCACGTGACCTTCCATCCGGCAATGATGCCGTTGACGCGTTCGATA |
| Mutant 10 | F27L_N73D_V103I | TCGGCTAGCGAGGTCCAACTTCTAGAATCTGGTGGGGGGCTTGTCCAGCCTGGTGGCACTTTGAGGCTGTCTTGTGCTGCCTCAGGGT<br>TGATCGTAAGTAGTAATTACATGAGTTGGGTGCGTCAGGCACCCGGAAAAGGGCTTGAATGGGTTAGTTTAGTTTACCCAGGTGGTAGCA<br>CTTACTATGCCGACTCTGTCAAAGGTAGGTTCACCGTTAGTCGTGATGATTCTAAAAACACTTTATACCTGCAAATGAATTCACTACGTGCT<br>GAAGACATGGCTGTTTATTATTGCGCACGTGACCTTCCATCCGGCATTGATGCCGTTGACGCGTTCGATA |
| Mutant 11 | F27L_V29A_N73D_V103I | TCGGCTAGCGAGGTCCAACTTCTAGAATCTGGTGGGGGGCTTGTCCAGCCTGGTGGCACTTTGAGGCTGTCTTGTGCTGCCTCAGGGT<br>TGATCGCTAGTAGTAATTACATGAGTTGGGTGCGTCAGGCACCCGGAAAAGGGCTTGAATGGGTTAGTTTAGTTTACCCAGGTGGTAGCA<br>CTTACTATGCCGACTCTGTCAAAGGTAGGTTCACCGTTAGTCGTGATGATTCTAAAAACACTTTATACCTGCAAATGAATTCACTACGTGCT<br>GAAGACATGGCTGTTTATTATTGCGCACGTGACCTTCCATCCGGCATTGATGCCGTTGACGCGTTCGATA |
| Mutant 12 | F27L_V29A_V103I | TCGGCTAGCGAGGTCCAACTTCTAGAATCTGGTGGGGGGCTTGTCCAGCCTGGTGGCACTTTGAGGCTGTCTTGTGCTGCCTCAGGGT<br>TGATCGCTAGTAGTAATTACATGAGTTGGGTGCGTCAGGCACCCGGAAAAGGGCTTGAATGGGTTAGTTTAGTTTACCCAGGTGGTAGCA<br>CTTACTATGCCGACTCTGTCAAAGGTAGGTTCACCGTTAGTCGTGATAATTCTAAAAACACTTTATACCTGCAAATGAATTCACTACGTGCT<br>GAAGACATGGCTGTTTATTATTGCGCACGTGACCTTCCATCCGGCATTGATGCCGTTGACGCGTTCGATA |
| Mutant 13 | F27L_V29D_V103I | TCGGCTAGCGAGGTCCAACTTCTAGAATCTGGTGGGGGGCTTGTCCAGCCTGGTGGCACTTTGAGGCTGTCTTGTGCTGCCTCAGGGT<br>TGATCGATAGTAGTAATTACATGAGTTGGGTGCGTCAGGCACCCGGAAAAGGGCTTGAATGGGTTAGTTTAGTTTACCCAGGTGGTAGCA<br>CTTACTATGCCGACTCTGTCAAAGGTAGGTTCACCGTTAGTCGTGATAATTCTAAAAACACTTTATACCTGCAAATGAATTCACTACGTGCT<br>GAAGACATGGCTGTTTATTATTGCGCACGTGACCTTCCATCCGGCATTGATGCCGTTGACGCGTTCGATA |

| | | |
|---|---|---|
| Mutant 14 | F27L_V29E_V103I | TCGGCTAGCGAGGTCCAACTTCTAGAATCTGGTGGGGGGCTTGTCCAGCCTGGTGGCACTTTGAGGCTGTCTTGTGCTGCCTCAGGGT<br>TGATCGAAAGTAGTAATTACATGAGTTGGGTGCGTCAGGCACCCGGAAAAGGGCTTGAATGGGTTAGTTTAGTTTACCCAGGTGGTAGCA<br>CTTACTATGCCGACTCTGTCAAAGGTAGGTTCACCGTTAGTCGTGATAATTCTAAAAACACTTTATACCTGCAAATGAATTCACTACGTGCT<br>GAAGACATGGCTGTTTATTATTGCGCACGTGACCTTCCATCCGGCATTGATGCCGTTGACGCGTTCGATA |
| Mutant 15 | G26E_F27L_N73D_V103I | TCGGCTAGCGAGGTCCAACTTCTAGAATCTGGTGGGGGGCTTGTCCAGCCTGGTGGCACTTTGAGGCTGTCTTGTGCTGCCTCAGAATT<br>GATCGTAAGTAGTAATTACATGAGTTGGGTGCGTCAGGCACCCGGAAAAGGGCTTGAATGGGTTAGTTTAGTTTACCCAGGTGGTAGCAC<br>TTACTATGCCGACTCTGTCAAAGGTAGGTTCACCGTTAGTCGTGATGATTCTAAAAACACTTTATACCTGCAAATGAATTCACTACGTGCTG<br>AAGACATGGCTGTTTATTATTGCGCACGTGACCTTCCATCCGGCATTGATGCCGTTGACGCGTTCGATA |
| Mutant 16 | G26E_F27L_N73D | TCGGCTAGCGAGGTCCAACTTCTAGAATCTGGTGGGGGGCTTGTCCAGCCTGGTGGCACTTTGAGGCTGTCTTGTGCTGCCTCAGAATT<br>GATCGTAAGTAGTAATTACATGAGTTGGGTGCGTCAGGCACCCGGAAAAGGGCTTGAATGGGTTAGTTTAGTTTACCCAGGTGGTAGCAC<br>TTACTATGCCGACTCTGTCAAAGGTAGGTTCACCGTTAGTCGTGATGATTCTAAAAACACTTTATACCTGCAAATGAATTCACTACGTGCTG<br>AAGACATGGCTGTTTATTATTGCGCACGTGACCTTCCATCCGGCGTTGATGCCGTTGACGCGTTCGATA |
| Mutant 17 | G26E_F27I_N73I | TCGGCTAGCGAGGTCCAACTTCTAGAATCTGGTGGGGGGCTTGTCCAGCCTGGTGGCACTTTGAGGCTGTCTTGTGCTGCCTCAGAAAT<br>TATCGTAAGTAGTAATTACATGAGTTGGGTGCGTCAGGCACCCGGAAAAGGGCTTGAATGGGTTAGTTTAGTTTACCCAGGTGGTAGCAC<br>TTACTATGCCGACTCTGTCAAAGGTAGGTTCACCGTTAGTCGTGATATTTCTAAAAACACTTTATACCTGCAAATGAATTCACTACGTGCTG<br>AAGACATGGCTGTTTATTATTGCGCACGTGACCTTCCATCCGGCGTTGATGCCGTTGACGCGTTCGATA |
| Mutant 18 | G26E_F27I_N73S | TCGGCTAGCGAGGTCCAACTTCTAGAATCTGGTGGGGGGCTTGTCCAGCCTGGTGGCACTTTGAGGCTGTCTTGTGCTGCCTCAGAAAT<br>TATCGTAAGTAGTAATTACATGAGTTGGGTGCGTCAGGCACCCGGAAAAGGGCTTGAATGGGTTAGTTTAGTTTACCCAGGTGGTAGCAC<br>TTACTATGCCGACTCTGTCAAAGGTAGGTTCACCGTTAGTCGTGATTCTTCTAAAAACACTTTATACCTGCAAATGAATTCACTACGTGCTG<br>AAGACATGGCTGTTTATTATTGCGCACGTGACCTTCCATCCGGCGTTGATGCCGTTGACGCGTTCGATA |
| Mutant 19 | G26E_F27L_N73S_V103S | TCGGCTAGCGAGGTCCAACTTCTAGAATCTGGTGGGGGGCTTGTCCAGCCTGGTGGCACTTTGAGGCTGTCTTGTGCTGCCTCAGAATT<br>GATCGTAAGTAGTAATTACATGAGTTGGGTGCGTCAGGCACCCGGAAAAGGGCTTGAATGGGTTAGTTTAGTTTACCCAGGTGGTAGCAC<br>TTACTATGCCGACTCTGTCAAAGGTAGGTTCACCGTTAGTCGTGATTCTTCTAAAAACACTTTATACCTGCAAATGAATTCACTACGTGCTG<br>AAGACATGGCTGTTTATTATTGCGCACGTGACCTTCCATCCGGCTCTGATGCCGTTGACGCGTTCGATA |
| Mutant 20 | G26E_F27L_N73S | TCGGCTAGCGAGGTCCAACTTCTAGAATCTGGTGGGGGGCTTGTCCAGCCTGGTGGCACTTTGAGGCTGTCTTGTGCTGCCTCAGAATT<br>GATCGTAAGTAGTAATTACATGAGTTGGGTGCGTCAGGCACCCGGAAAAGGGCTTGAATGGGTTAGTTTAGTTTACCCAGGTGGTAGCAC<br>TTACTATGCCGACTCTGTCAAAGGTAGGTTCACCGTTAGTCGTGATTCTTCTAAAAACACTTTATACCTGCAAATGAATTCACTACGTGCTG<br>AAGACATGGCTGTTTATTATTGCGCACGTGACCTTCCATCCGGCGTTGATGCCGTTGACGCGTTCGATA |
| Mutant 21 | G26E_F27I | TCGGCTAGCGAGGTCCAACTTCTAGAATCTGGTGGGGGGCTTGTCCAGCCTGGTGGCACTTTGAGGCTGTCTTGTGCTGCCTCAGAAAT<br>TATCGTAAGTAGTAATTACATGAGTTGGGTGCGTCAGGCACCCGGAAAAGGGCTTGAATGGGTTAGTTTAGTTTACCCAGGTGGTAGCAC<br>TTACTATGCCGACTCTGTCAAAGGTAGGTTCACCGTTAGTCGTGATAATTCTAAAAACACTTTATACCTGCAAATGAATTCACTACGTGCTG<br>AAGACATGGCTGTTTATTATTGCGCACGTGACCTTCCATCCGGCGTTGATGCCGTTGACGCGTTCGATA |
| Mutant 22 | G26E_F27L | TCGGCTAGCGAGGTCCAACTTCTAGAATCTGGTGGGGGGCTTGTCCAGCCTGGTGGCACTTTGAGGCTGTCTTGTGCTGCCTCAGAATT<br>GATCGTAAGTAGTAATTACATGAGTTGGGTGCGTCAGGCACCCGGAAAAGGGCTTGAATGGGTTAGTTTAGTTTACCCAGGTGGTAGCAC<br>TTACTATGCCGACTCTGTCAAAGGTAGGTTCACCGTTAGTCGTGATAATTCTAAAAACACTTTATACCTGCAAATGAATTCACTACGTGCTG<br>AAGACATGGCTGTTTATTATTGCGCACGTGACCTTCCATCCGGCGTTGATGCCGTTGACGCGTTCGATA |
| Mutant 23 | G26E_F27I_N73T | TCGGCTAGCGAGGTCCAACTTCTAGAATCTGGTGGGGGGCTTGTCCAGCCTGGTGGCACTTTGAGGCTGTCTTGTGCTGCCTCAGAAAT<br>TATCGTAAGTAGTAATTACATGAGTTGGGTGCGTCAGGCACCCGGAAAAGGGCTTGAATGGGTTAGTTTAGTTTACCCAGGTGGTAGCAC<br>TTACTATGCCGACTCTGTCAAAGGTAGGTTCACCGTTAGTCGTGATACTTCTAAAAACACTTTATACCTGCAAATGAATTCACTACGTGCTG<br>AAGACATGGCTGTTTATTATTGCGCACGTGACCTTCCATCCGGCGTTGATGCCGTTGACGCGTTCGATA |
| Mutant 24 | F27I_N73D_V103I | TCGGCTAGCGAGGTCCAACTTCTAGAATCTGGTGGGGGGCTTGTCCAGCCTGGTGGCACTTTGAGGCTGTCTTGTGCTGCCTCAGGGA<br>TTATCGTAAGTAGTAATTACATGAGTTGGGTGCGTCAGGCACCCGGAAAAGGGCTTGAATGGGTTAGTTTAGTTTACCCAGGTGGTAGCA<br>CTTACTATGCCGACTCTGTCAAAGGTAGGTTCACCGTTAGTCGTGATGATTCTAAAAACACTTTATACCTGCAAATGAATTCACTACGTGCT<br>GAAGACATGGCTGTTTATTATTGCGCACGTGACCTTCCATCCGGCATTGATGCCGTTGACGCGTTCGATA |
| Mutant 25 | G26E_F27L_V103I | TCGGCTAGCGAGGTCCAACTTCTAGAATCTGGTGGGGGGCTTGTCCAGCCTGGTGGCACTTTGAGGCTGTCTTGTGCTGCCTCAGAATT<br>GATCGTAAGTAGTAATTACATGAGTTGGGTGCGTCAGGCACCCGGAAAAGGGCTTGAATGGGTTAGTTTAGTTTACCCAGGTGGTAGCAC<br>TTACTATGCCGACTCTGTCAAAGGTAGGTTCACCGTTAGTCGTGATAATTCTAAAAACACTTTATACCTGCAAATGAATTCACTACGTGCTG<br>AAGACATGGCTGTTTATTATTGCGCACGTGACCTTCCATCCGGCATTGATGCCGTTGACGCGTTCGATA |
| Mutant 26 (Delta-63) | G26E_F27L_N73E_V103I | tcggctagcgaggtccaacttctagaatctggtggggggcttgtccagcctggtggcactttgaggctgtcttgtgctgcctcagaattgatcgtaagtagtaattacatg<br>agttgggtgcgtcaggcacccggaaaagggcttgaatgggttagtttagtttacccaggtggtagcacttactatgccgactctgtcaaaggtaggttcaccgttagtcg<br>tgatgaatctaaaaacactttatacctgcaaatgaattcactacgtgctgaagacatggctgtttattattgcgcacgtgaccttccatccggcattgatgccgttgacgc<br>gttcgata |
| Mutant 27 | F27L_N73I_V103I | TCGGCTAGCGAGGTCCAACTTCTAGAATCTGGTGGGGGGCTTGTCCAGCCTGGTGGCACTTTGAGGCTGTCTTGTGCTGCCTCAGGGT<br>TGATCGTAAGTAGTAATTACATGAGTTGGGTGCGTCAGGCACCCGGAAAAGGGCTTGAATGGGTTAGTTTAGTTTACCCAGGTGGTAGCA<br>CTTACTATGCCGACTCTGTCAAAGGTAGGTTCACCGTTAGTCGTGATATTTCTAAAAACACTTTATACCTGCAAATGAATTCACTACGTGCT<br>GAAGACATGGCTGTTTATTATTGCGCACGTGACCTTCCATCCGGCATTGATGCCGTTGACGCGTTCGATA |
| Mutant 28 | F27I_N73I_V103I | TCGGCTAGCGAGGTCCAACTTCTAGAATCTGGTGGGGGGCTTGTCCAGCCTGGTGGCACTTTGAGGCTGTCTTGTGCTGCCTCAGGGA<br>TTATCGTAAGTAGTAATTACATGAGTTGGGTGCGTCAGGCACCCGGAAAAGGGCTTGAATGGGTTAGTTTAGTTTACCCAGGTGGTAGCA<br>CTTACTATGCCGACTCTGTCAAAGGTAGGTTCACCGTTAGTCGTGATATTTCTAAAAACACTTTATACCTGCAAATGAATTCACTACGTGCT<br>GAAGACATGGCTGTTTATTATTGCGCACGTGACCTTCCATCCGGCATTGATGCCGTTGACGCGTTCGATA |

**SUPPLEMENTARY FIGURES**

**Figure S1:** Performance of different model architectures on held-out test data for the Absolut! synthetic data experiments

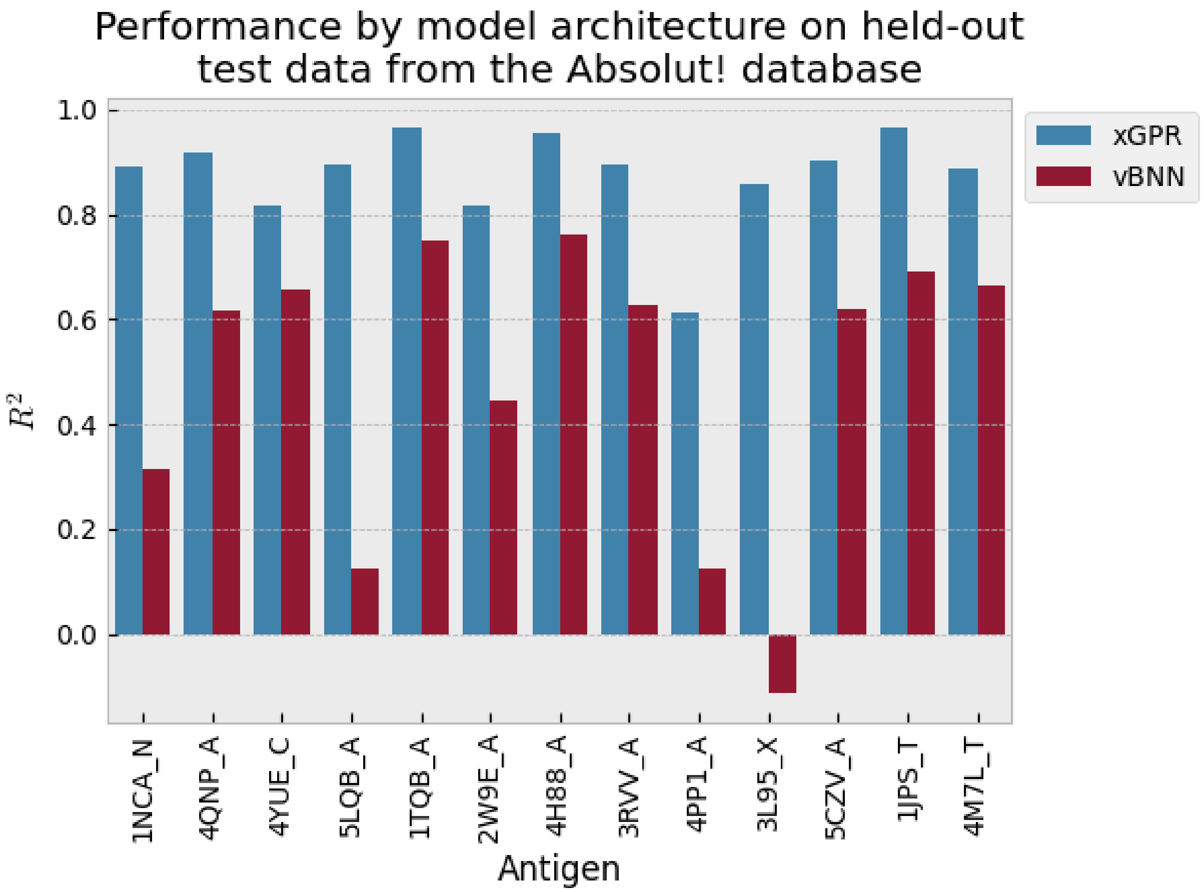

**Figure S2:** Sequence alignment of the Delta-6 and Delta-63 scFv discovered in this work. Alignment performed with Expasy SIM alignment tool. Mutated residues in bold.

Delta-6: EVQLLESGGGLVQPGGTLRLSCAASGFIVSSNYMSWVRQAPGKGLEWVSLVYPGGSTYYA

Delta63: EVQLLESGGGLVQPGGTLRLSCAAS**EL**IVSSNYMSWVRQAPGKGLEWVSLVYPGGSTYYA

************************* *********************************

Delta-6: DSVKGRFTVSRDNSKNTLYLQMNSLRAEDMAVYYCARDLPSGVDAVDAFDIWGQGTMVTV

Delta63: DSVKGRFTVSRD**E**SKNTLYLQMNSLRAEDMAVYYCARDLPSG**I**DAVDAFDIWGQGTMVTV

************ ***************************** *****************

Delta-6: SSGILGSGGGGSGGGGSGGGGSDIRVTQSPSSLSASVGDRVSITCRASQIISGYLNWYQQ

```
Delta63:  SSGILGSGGGGSGGGGSGGGGSDIRVTQSPSSLSASVGDRVSITCRASQIISGYLNWYQQ

          ************************************************************

Delta-6:   KPGSAPQLLIYASSSLQSGVPPRFSGSRSGTEFTLTISSLQPEDFATYYCQQTYSIPFTF

Delta63:   KPGSAPQLLIYASSSLQSGVPPRFSGSRSGTEFTLTISSLQPEDFATYYCQQTYSIPFTF

           ************************************************************

Delta-6:   GPGTKVDIK

Delta63:   GPGTKVDIK

           *********
```

**Figure S3:** Addition of a large molar excess of ACE2 receptor reduces Delta-63 scFv binding to the SARS COV-2 WT RBD on the yeast surface. MFI is mean fluorescent intensity of binding.

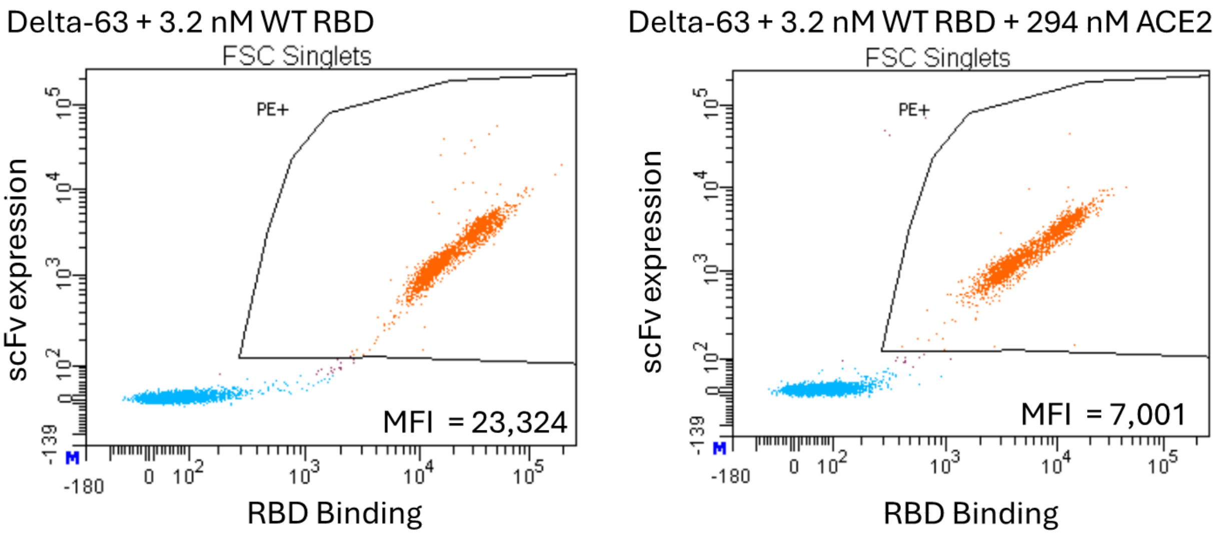

**Figure S4:** Expression level of (A) Delta-6 vs. Delta-63 on the yeast surface and (B) Delta-63 vs. anti-COVID or anti-PDL1 scFv from EY6A[1], CR3022[2], S103F/S33R CR3022[3], Beta-54[4], S304[5], and Atezolizumab[6] on the yeast surface. Expression levels were measured by expression of the c-terminal c-myc tag of each construct via flow cytometry in replicates. MFI is mean fluorescent intensity.

(A)

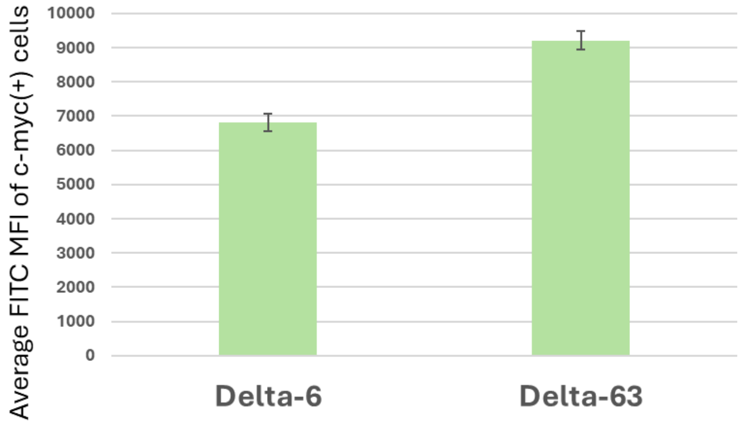

(B)

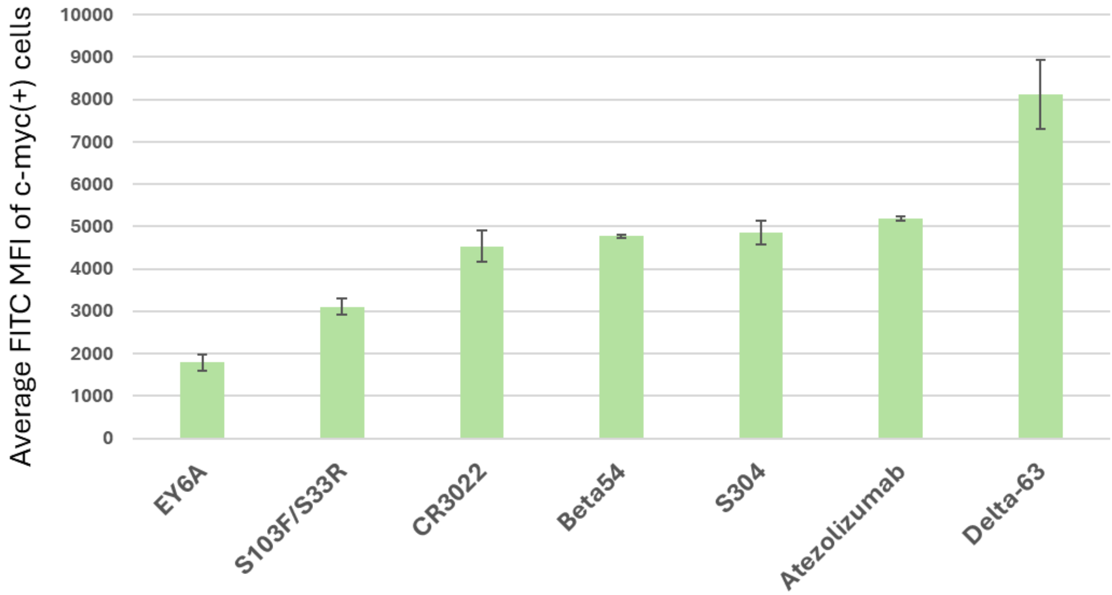

Average FITC MFI of c-myc(+) cells
10000
9000
8000
7000
6000
5000
4000
3000
2000
1000
0
EY6A
S103F/S33R
CR3022
Beta54
S304
Atezolizumab
Delta-63

**Figure S5: Example FACS plots of Delta-6 library against different RBD antigens.** (A) RBDs where the $K_D$ value for WT Delta-6 scFv were in the low nM range, requiring only a single sort (cells in “Hi” gate were better than WT Delta-6 scFv, while cells in the blue gate were approximately WT level $K_D$ binders). (B) Example of an RBD which showed no binding to the WT Delta-6 scFv, requiring two sorts to enrich moderate (130 nM) binders or tighter (30 nM) binders (collected cells were from “Hi” gates). The binders collected under the 130 nM and 30 nM conditions were given unique indices to distinguish them in the MiSeq sequencing run (as were all cells collected under different antigen concentrations). For all plots, the y-axis represents scFv expression intensity while x-axis is RBD binding intensity.

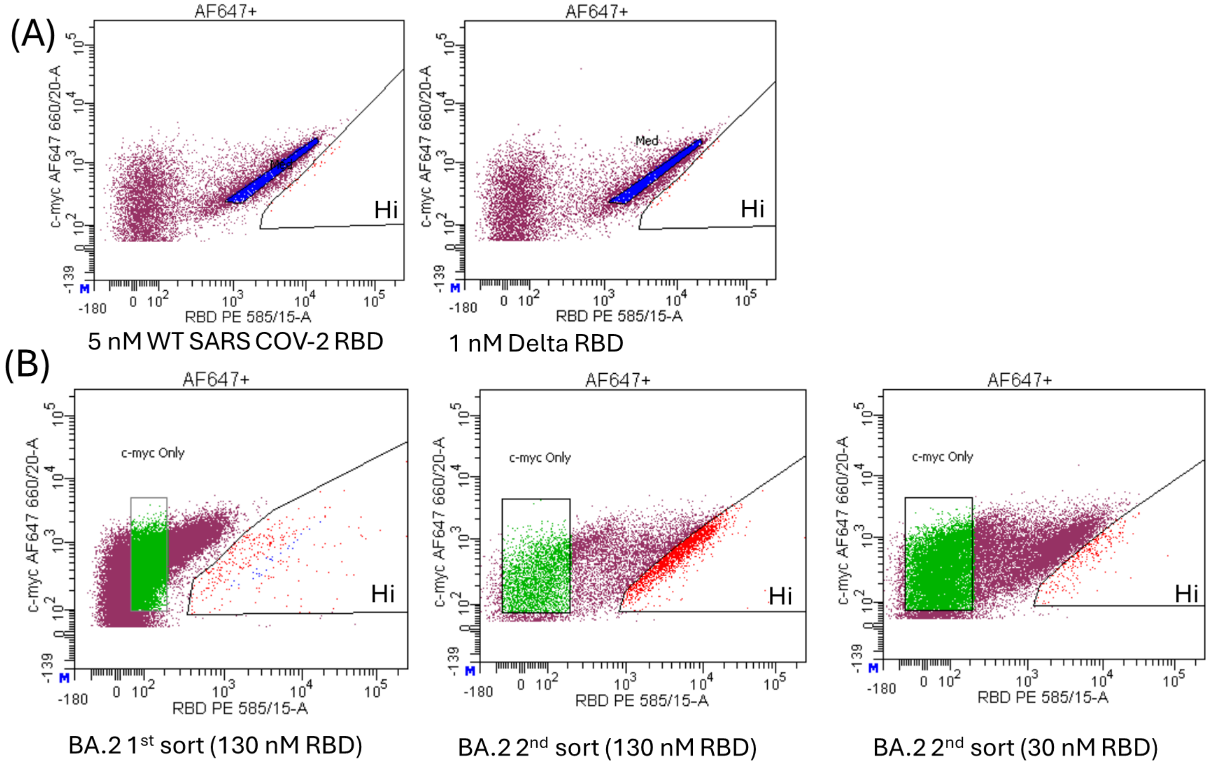

**Figure S6.** WT Delta-6 scFv in pCTCON2 (nucleotide sequence), bold region is scFv sequence while edges (non bold font) are from the pCTCON2 vector.

Gcggtagcggaggcggagggtcggctagc**gaggtccaacttctagaatctggtggggggcttgtccagcctggtggc
actttgaggctgtcttgtgctgcctcagggttcatcgtaagtagtaattacatgagttgggtgcgtcaggcacccg
gaaaagggcttgaatgggttagtttagtttacccaggtggtagcacttactatgccgactctgtcaaaggtaggttc
accgttagtcgtgataattctaaaaacactttatacctgcaaatgaattcactacgtgctgaagacatggctgtttatt
attgcgcacgtgaccttccatccggcgttgatgccgttgacgcgttcgatatatggggccaaggtactatggtga
cggtcagttcagggattctaggaagtggaggtgggggtccggcggcggaggctcaggagggggcggttct
gacataagagttacccagagcccgagttctctgtctgcatctgtgggtgacagagtttcaatcacgtgcagagca
tctcaaataatctccggttatctaaactggtaccagcaaaagcccggatctgcaccccagttattaatttacgcaa
gtagctcccttcaaagtggcgttccgccacgtttttctggtagccgtagtgggaccgaatttacattgacaatctca
tccctgcagccggaggacttcgcaacgtattactgccaacagacgtactcaattccgtttacttttggccgggc
acaaaggtcgatattaaa**ggtggcggatccgaacaaaagcttatttctgaagaggacttgtaa

**Figure S7**. Flow cytometry analysis of the WT Delta-6 scFv vs. 29-member library using the 10 RBDs used for the Delta-6 library sorts. The % values represent % of expressing library members above the WT Delta-6 main population. As a control, the % of WT Delta-6 cells appearing above the WT main group is included. The y-axis of each plot represents scFv expression (c-myc tag) while the x-axis the binding intensity to the RBD (PE fluorescence). Also included are plots of the WT Delta-6 scFv and 29-member library with no RBD (just streptavidin-PE reagent). The WT/29-member library vs. WT SARS COV-2 RBD includes a replicate measurement.

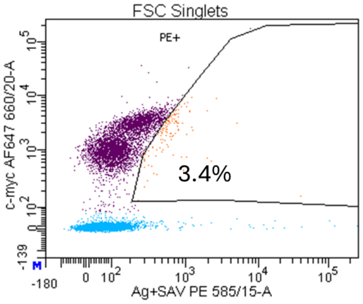

Delta-6 Beta RBD 321 nM

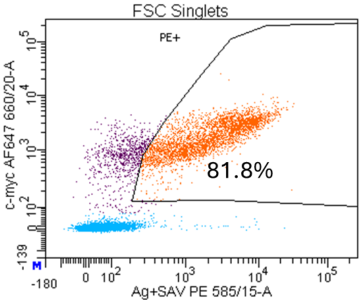

29-member library Beta RBD 321 nM

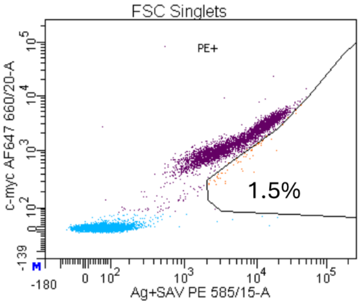

Delta-6 Alpha RBD 6 nM

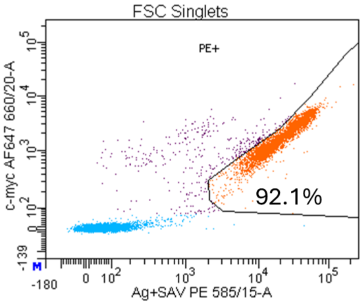

29-member library Alpha RBD 6 nM

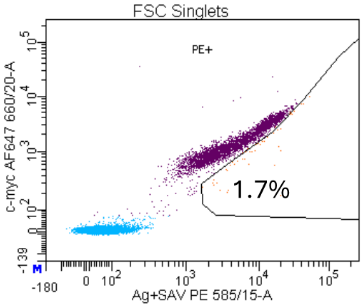

Delta-6 WT RBD 6.3 nM

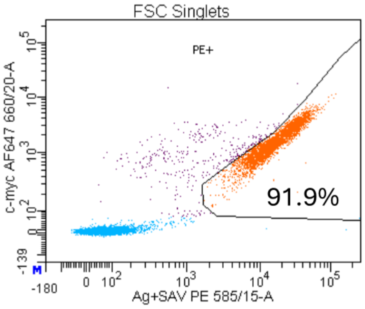

29-member library WT RBD 6.3 nM

Replicate 1

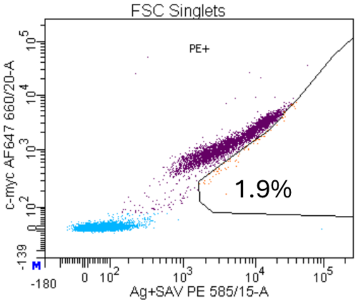

Delta-6 WT RBD 6.3 nM

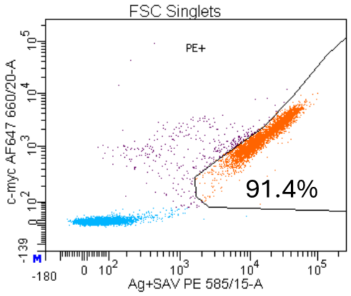

29-member library WT RBD 6.3 nM

Replicate 2

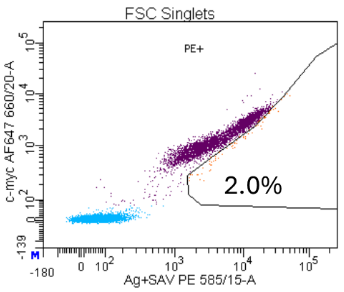

Delta-6 Lambda RBD 2.6 nM

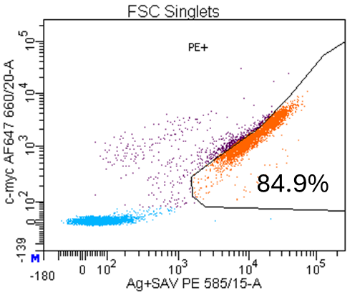

29-member library Lambda RBD 2.6 nM

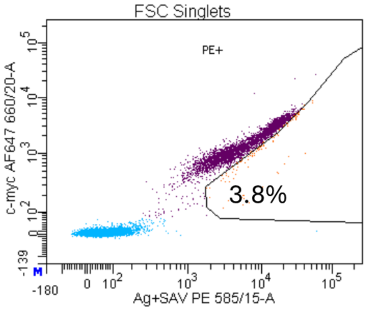

Delta-6 Delta RBD 1.3 nM

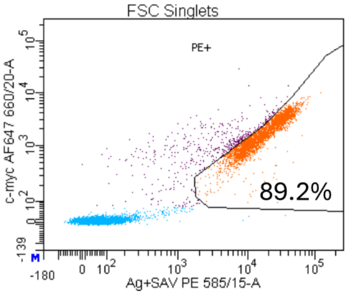

29-member library Delta RBD 1.3 nM

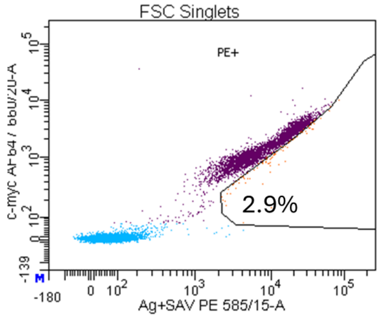

Delta-6 Kappa RBD 160 nM

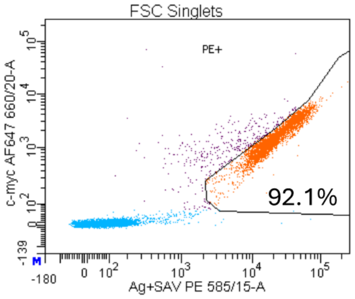

29-member library Kappa RBD 160 nM

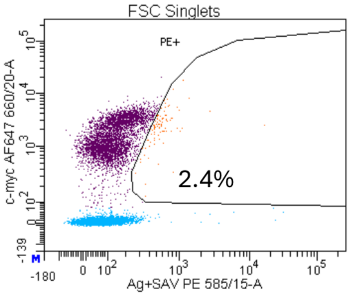

Delta-6 Gamma RBD 321 nM

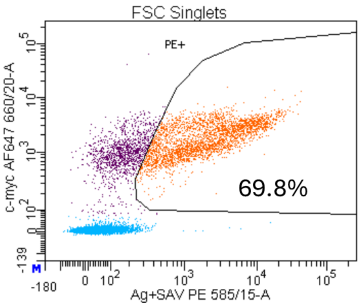

29-member library Gamma RBD 321 nM

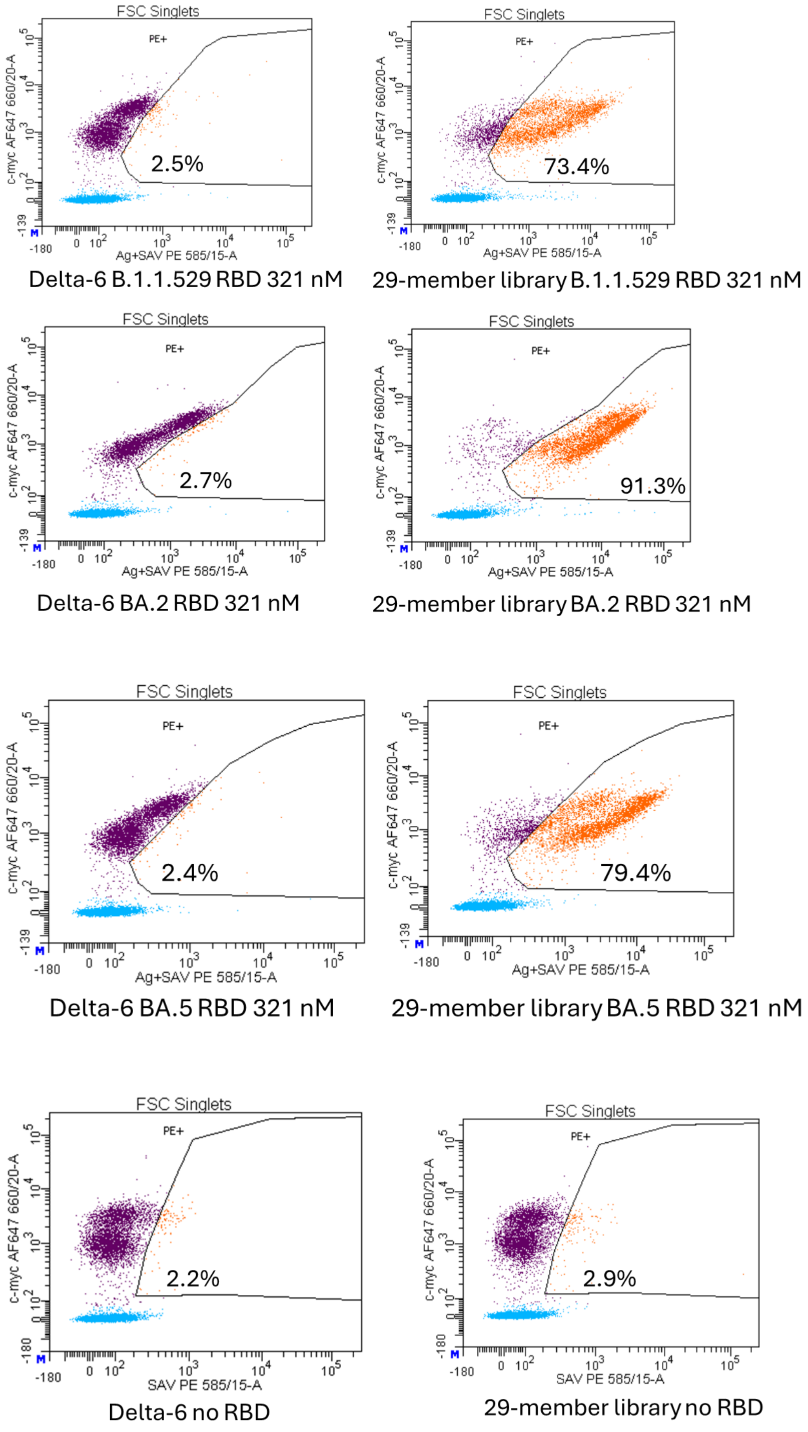

**Figure S8**. Binding curves generated by yeast surface display against 10 RBDs for both the WT Delta-6 and mutant Delta-63 scFv. Y-axis is phycoerythrin (PE) mean fluorescent intensity (MFI) of yeast, while x-axis is concentration of titrated RBD in nM. For RBDs which bound too weakly to determine a $K_D$, the data points are displayed in Excel format. The rest of the curves were plotted with GraphPad Prism 10 software.

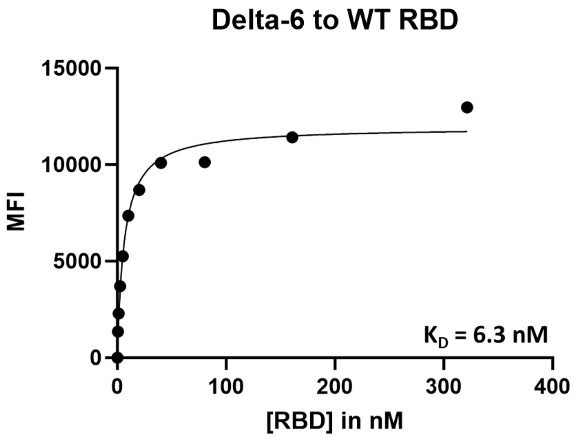
Delta-6 to WT RBD
15000
10000
5000
0
MFI
K_D = 6.3 nM
0
100
200
300
400
[RBD] in nM

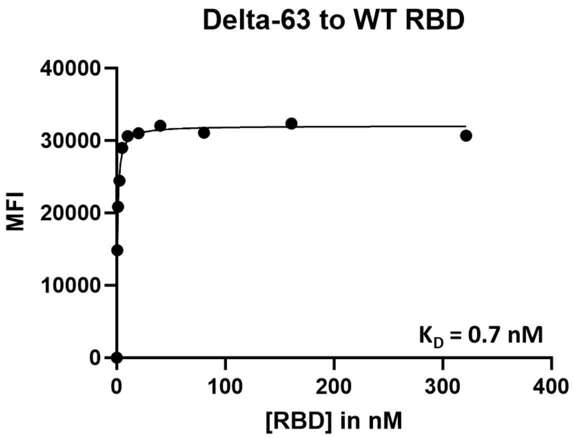
Delta-63 to WT RBD
40000
30000
20000
10000
0
MFI
K_D = 0.7 nM
0
100
200
300
400
[RBD] in nM

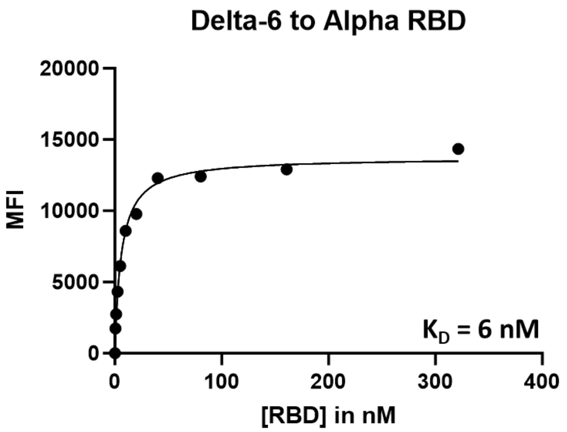
Delta-6 to Alpha RBD
20000
15000
10000
5000
0
MFI
K_D = 6 nM
0
100
200
300
400
[RBD] in nM

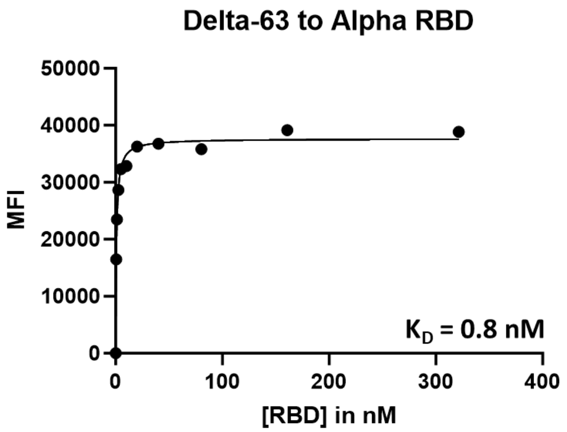
Delta-63 to Alpha RBD
50000
40000
30000
20000
10000
0
MFI
K_D = 0.8 nM
0
100
200
300
400
[RBD] in nM

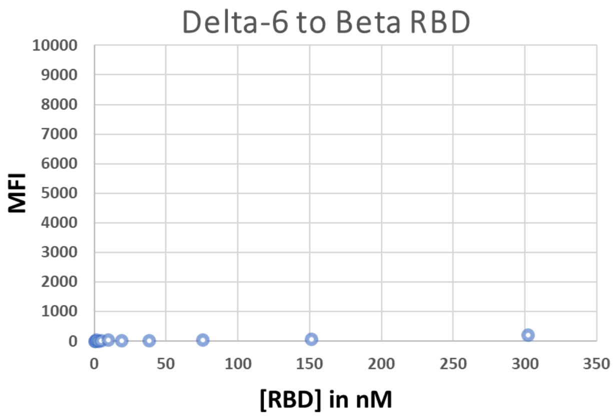
Delta-6 to Beta RBD
10000
9000
8000
7000
6000
5000
4000
3000
2000
1000
0
MFI
0
50
100
150
200
250
300
350
[RBD] in nM

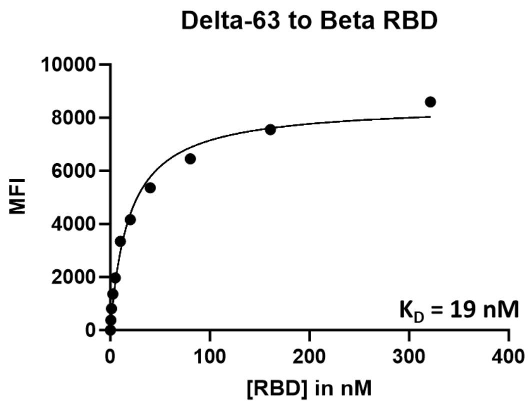
Delta-63 to Beta RBD
10000
8000
6000
4000
2000
0
MFI
K_D = 19 nM
0
100
200
300
400
[RBD] in nM

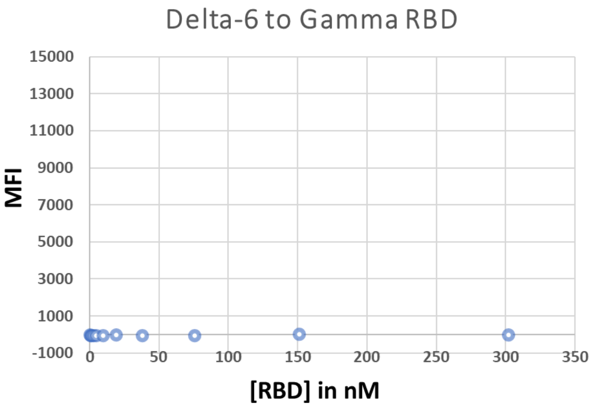
Delta-6 to Gamma RBD
MFI
15000
13000
11000
9000
7000
5000
3000
1000
-1000
0
50
100
150
200
250
300
350
[RBD] in nM

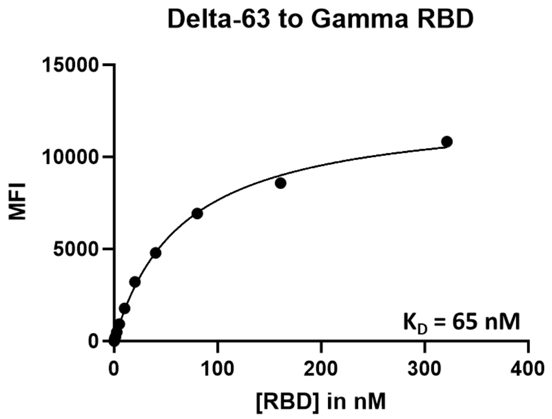
Delta-63 to Gamma RBD
MFI
15000
10000
5000
0
$K_D$ = 65 nM
0
100
200
300
400
[RBD] in nM

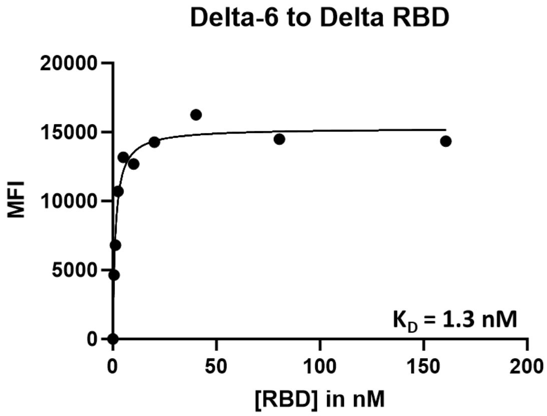
Delta-6 to Delta RBD
MFI
20000
15000
10000
5000
0
$K_D$ = 1.3 nM
0
50
100
150
200
[RBD] in nM

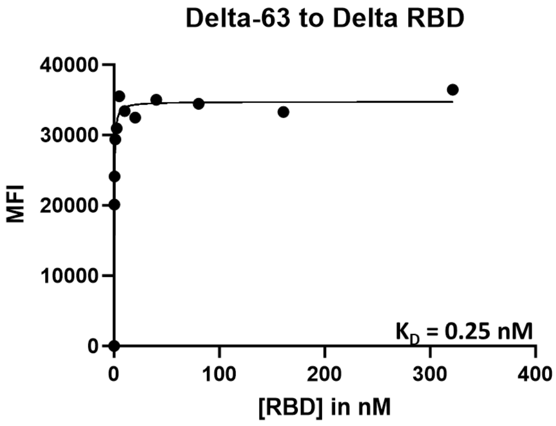
Delta-63 to Delta RBD
MFI
40000
30000
20000
10000
0
$K_D$ = 0.25 nM
0
100
200
300
400
[RBD] in nM

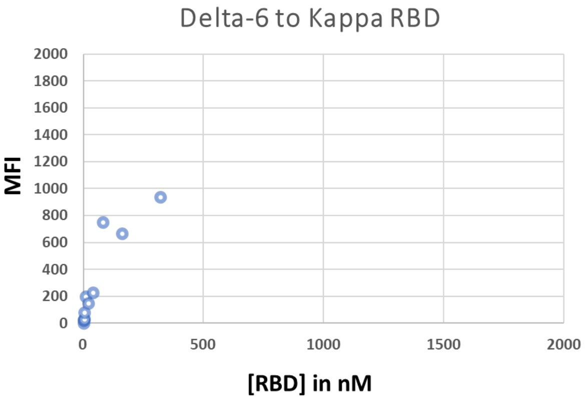
Delta-6 to Kappa RBD
MFI
2000
1800
1600
1400
1200
1000
800
600
400
200
0
0
500
1000
1500
2000
[RBD] in nM

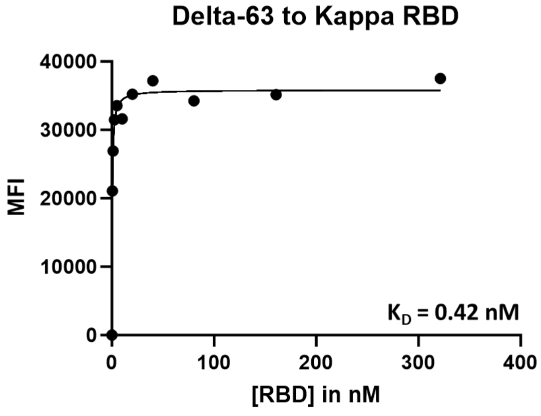
Delta-63 to Kappa RBD
MFI
40000
30000
20000
10000
0
$K_D$ = 0.42 nM
0
100
200
300
400
[RBD] in nM

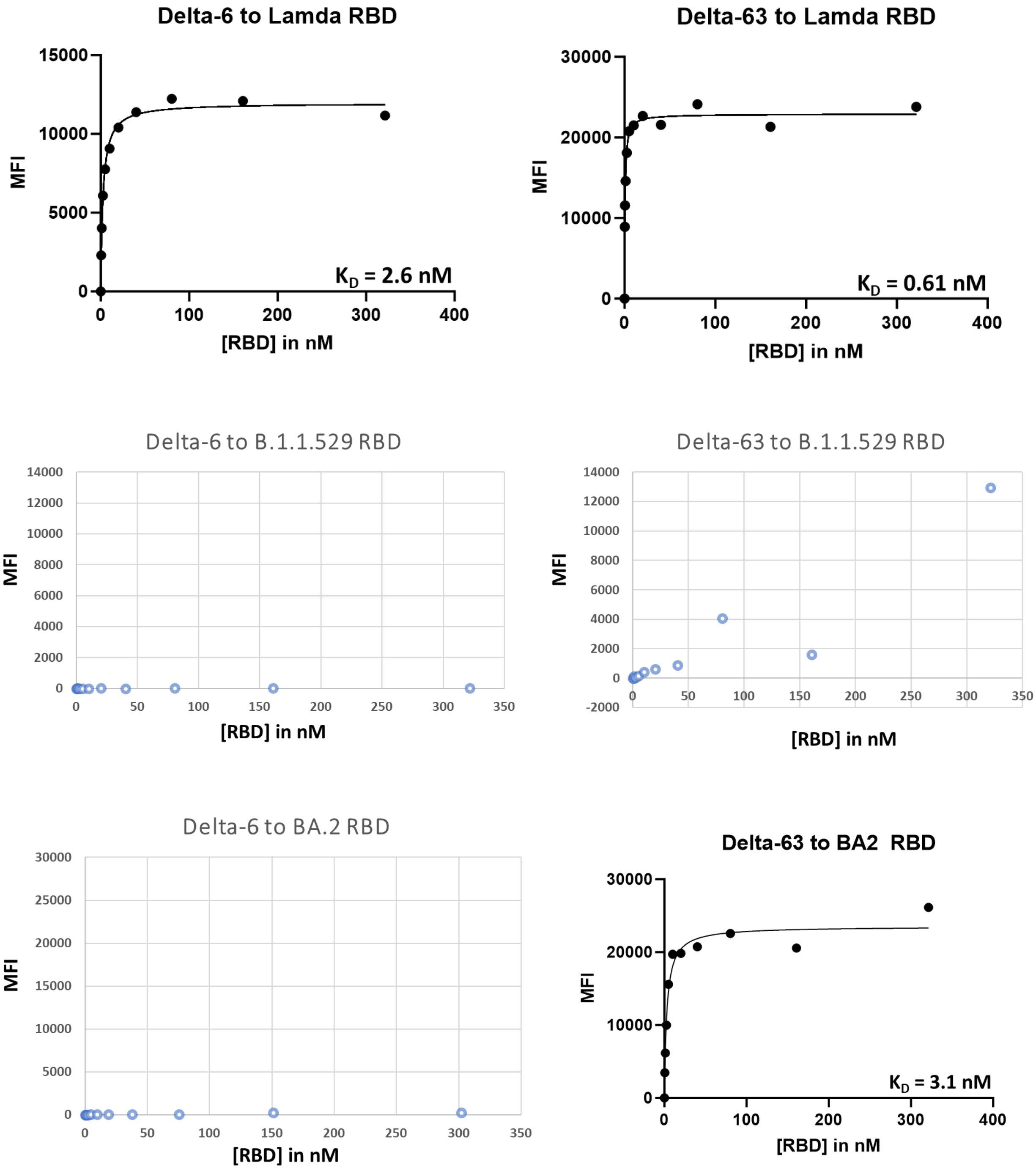

Delta-6 to Lamda RBD
MFI
[RBD] in nM
$K_D$ = 2.6 nM
Delta-63 to Lamda RBD
MFI
[RBD] in nM
$K_D$ = 0.61 nM
Delta-6 to B.1.1.529 RBD
MFI
[RBD] in nM
Delta-63 to B.1.1.529 RBD
MFI
[RBD] in nM
Delta-6 to BA.2 RBD
MFI
[RBD] in nM
Delta-63 to BA2 RBD
MFI
[RBD] in nM
$K_D$ = 3.1 nM

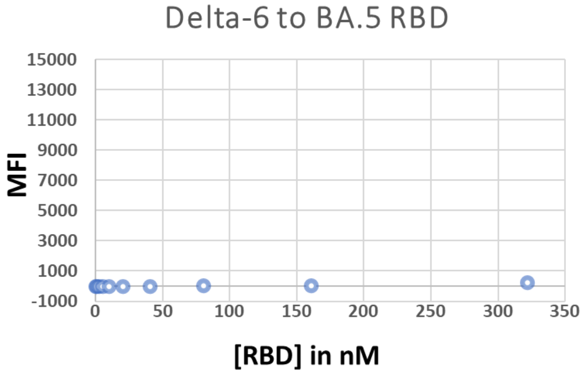

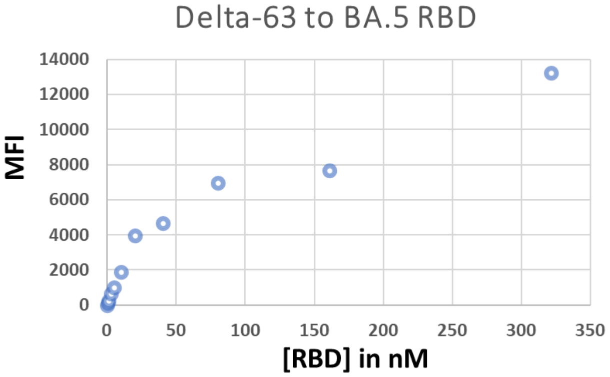
